# Zebrafish models of Mucopolysaccharidosis IIIA, IIIB, and IIIC show hyperactivity, neuroinflammation and changes in oligodendrocyte cell state

**DOI:** 10.1101/2023.08.02.550904

**Authors:** Ewan Gerken, Syahida Ahmad, Lakshay Rattan, Kim Hemsley, Angel Allen, Adeline Lau, Karissa Barthelson, Michael Lardelli

## Abstract

Sanfilippo syndrome, also known as mucopolysaccharidosis type III (MPS III), is a group of rare inherited lysosomal storage disorders which causes childhood dementia. Subtypes of MPS III are caused by deficiencies in one of four enzymes required for degradation of the glycosaminoglycan heparan sulfate (HS). An inability to degrade HS leads to progressive neurodegeneration and death, often in the second or third decades of life in the classical course of the disease. Knowledge of MPS III pathogenesis is incomplete, and no effective therapies are yet approved for human use. We generated hypomorphic mutations in the endogenous zebrafish genes orthologous to those associated with MPS IIIA, MPS IIIB and MPS IIIC. Our models display the primary MPS III disease signature of significant brain accumulation of HS, and behavioural analyses revealed hyperactivity phenotypes in MPS IIIA and MPS IIIB zebrafish. Brain transcriptome analysis revealed changes related to lysosomal, glycosaminoglycan and immune system biology in all three models but also distinct differences in brain transcriptome state between models, which may be mutation dependent. The transcriptome analysis also indicated marked disturbance of oligodendrocyte-related genes in the brains of MPS III zebrafish, supporting that effects on this cell type are a consistent characteristic of MPS III. Overall, our zebrafish models recapture key characteristics of the human disease and phenotypes seen in mouse models. Our models will allow exploitation of the zebrafish’s extreme fecundity and accessible anatomy to dissect the pathological mechanisms in MPS IIIA, IIIB, and IIIC.

## Introduction

Sanfilippo syndrome, also known as Mucopolysaccharidosis type III (MPS III) is caused by rare, autosomal recessive lysosomal storage disorders that result in childhood dementia. MPS III results from mutations in one of four genes encoding enzymes involved in the degradation of the glycosaminoglycan (GAG) heparan sulfate (HS), with four subtypes A, B, C, and D, distinguished by mutations in the *N*-sulfoglucosamine sulfohydrolase (*SGSH*), *N*-acetyl-alpha-glucosaminidase (*NAGLU*), heparan-alpha-glucosaminide *N*-acetyltransferase (*HGSNAT*), and glucosamine (*N*-acetyl)-6-sulfatase (*GNS*) genes respectively. Deficiency in any one of these enzymes leads to accumulation of incompletely degraded HS inside cells and in bodily fluids.

MPS III presents enormous challenges to the families of affected children due to the broad and progressive cognitive and behavioural changes that accompanying the disease. Early in the disease course, children are slow or fail to achieve normal developmental milestones. This is commonly followed by the onset of behavioural symptoms such as hyperactivity, aggression and sleep disturbances (Cleary and Wraith, 1993; Muschol et al., 2022; Valstar et al., 2011). Later stages of the disease involve progressive loss of cognition, speech, mobility and, eventually, deterioration of autonomic bodily functions such as swallowing and respiration (Valstar et al., 2011; Valstar et al., 2010). Whilst the incidence varies across geographic regions (Zelei et al., 2018), MPS IIIA is generally both the most common and, usually, most rapidly-progressing subtype, with median age at death estimated at 14.5 ± 4.2 years, followed by 15.4 ± 7.3 years for MPS IIIB and 22.4 ± 9.5 years for MPS IIIC (Lavery et al., 2017). A lack of data exists for MPS IIID due to its rarity. Significant heterogeneity also exists in the clinical presentation and rate of disease progression in MPS III, both between and within subtypes (Kamp et al., 1981), as well as between affected siblings (Andria et al., 1979; Lindor et al., 1994).

Analysis of post-mortem patient brain tissues has highlighted accumulation of incompletely degraded HS, gangliosides G_M2_ and G_M3_, misfolded proteins, along with significant neuroinflammation in several MPS disorders (Viana et al., 2020). However, animal models are required to dissect the precise interplay of the earlier molecular events leading to neurodegeneration in MPS III, which remain poorly understood. Identifying molecular and behavioural phenotypes that animal models share with human MPS III disease is important for screening and measuring the potential therapeutic benefit of pharmaceuticals. Currently, there are no approved curative treatments for MPS III, and the treatment options available are limited in effect and mostly supportive (Pearse and Iacovino, 2020). Therefore, there is a pressing need for novel, versatile animal models of the disease to advance our understanding of MPS III pathophysiology and for development of new therapies. While *SGSH, NAGLU* and *HGSNAT* encode enzymes in the same HS catabolic pathway, MPS III-causative mutations in these genes have never been compared in the closely related individuals of an animal model organism to see how their phenotypes may differ. Additionally, studies examining the differing sensitivities of the brain’s various cell types to MPS III-causative mutations are only recently emerging (Balak et al., 2024).

In recent years, the zebrafish (*Danio rerio*) has become a valuable model organism for studying lysosomal storage disorders. In total, over 60 zebrafish models of lysosomal storage disorders have been created and phenotyped (reviewed in (Zhang and Peterson, 2020)). Zebrafish show approximately 70% genetic similarity to humans (Herrero et al., 2016; Howe et al., 2013), with significant conservation observed across neurochemical systems and brain anatomy (Panula et al., 2010). Their small size, high fecundity, and rapid development make them an attractive system for high-throughput drug screening for discovery of therapeutics (reviewed in (Dash and Patnaik, 2023)). In addition, the transparency of zebrafish embryos and larvae enables easy visualisation and imaging of internal organs, tissues, and subcellular structures including lysosomes. In particular, the reproductive characteristics of zebrafish are advantageous for reducing genetic and environmental noise through intra-family comparisons between mutant and non-mutant genotypes. We have previously had success in this approach with transcriptome analyses of zebrafish carrying Alzheimer’s disease-like mutations (Barthelson et al., 2021; Barthelson, K. et al., 2020; Hin et al., 2020; Jiang et al., 2020). Zebrafish are also a suitable model for behavioural analysis, with evidence suggesting that regulatory mechanisms behind learning, memory, aggression, anxiety and sleep are conserved between zebrafish and mammals (reviewed in (Norton and Bally-Cuif, 2010)).

A zebrafish model of MPS IIIA has previously been described and recapitulates many features of the human disease including HS accumulation and behavioural deficits (Douek et al., 2021). Analysis of effects on brain proteome and transcriptome in this model revealed a variety of changes including to effectors of lysosomal catabolism and to immediate-early gene expression important in regulating postsynaptic neuronal responses (Douek et al., 2022). The brain transcriptome and proteome has also been described of a zebrafish model of MPS IIIB (Barthelson et al., 2025).

Here, we describe the creation and characterisation of the zebrafish model of MPS IIIB cited above, and novel models of MPS IIIA and MPS IIIC. We introduced endogenous mutations by CRISPR-Cas9 genome editing of the zebrafish genes *sgsh, naglu* and *hgsnat*. We excluded *gns* (MPS IIID) from our study due to this gene’s duplication in teleosts (Howe et al., 2013), complicating its suitability for modelling in zebrafish. We apply our previously successful intra-family strategy for comparison of different genotypes to investigate MPS III-driven changes in behaviour and brain transcriptome states. We use families of zebrafish siblings that include individuals of either of two MPS III subtypes to compare brain transcriptome effects between subtypes. Our results demonstrate that these zebrafish models recapitulate key features of MPS III, including GAG accumulation, metabolic defects, neuroinflammation, and behavioural deficits. Intriguingly, all three MPS III subtypes show disturbance of the oligodendrocyte cellular state suggesting that this neural cell type is particularly sensitive to loss of HS-degradative ability. These zebrafish MPS III models will serve as valuable tools for studying the pathophysiology of Sanfilippo syndrome and for the testing of novel therapeutic strategies.

## Methods

### Zebrafish husbandry and animal ethics statement

Experiments with zebrafish were performed under the auspices of the University of Adelaide Animal Ethics Committee (permit number S-2021-041) and Institutional Biosafety Committee (Dealing ID NLRD 15037). All experiments were performed with an inbred Tübingen (Tü) strain. Zebrafish embryos were collected and incubated in E3 embryo medium (Westerfield, 2000) at 28°C on a 14/10 hour light/dark cycle until 7 days post fertilisation (dpf). Hatched larvae were then moved to benchtop tanks (without water circulation) and fed live *Rotifera* from 7-21 dpf, followed by live *Artemia salina* from 21 dpf onwards. Juvenile fish were moved to recirculated water aquarium systems at 5 to 6 weeks post fertilisation. Adult zebrafish were maintained in shared recirculated water aquarium systems at 28°C at a density of 2-4 fish per litre (20-30 fish per 8L tank), on a 14/10 hour light/dark cycle, and fed NRD 5/8 dry flake food (Inve Aquaculture, Dendermonde, Belgium) in the morning and live *Artemia salina* (Inve Aquaculture) in the afternoon.

### Generation of hypomorphic mutations in zebrafish *sgsh, naglu* and *hgsnat*

#### Mutagenesis

Targeted knock-in mutations were generated in zebrafish using CRISPR-Cas9 (for *sgsh* and *hgsnat*) and CRISPR-Cpf1 (for *naglu*) systems (Integrated DNA Technologies, Coralville, IA, USA) via a non-homologous end joining approach. The sgRNAs used to target each gene are summarised in **Table S1**, and visualised in **Fig.S1**.

The sgRNAs for *sgsh* and *hgsnat* sgRNAs were co-injected into the same G_0_ embryos. For both genes, 1 μL of 100 μM crRNA (Integrated DNA Technologies) was mixed with 3 μL of tracrRNA (Integrated DNA Technologies) and 3 μL of nuclease-free duplex buffer (Integrated DNA Technologies). This solution was denatured at 95°C for 5 minutes, before being allowed to cool to room temperature to facilitate sgRNA heteroduplex formation. Then, 3 μL of 64 μM Cas9 3NLS nuclease (Integrated DNA Technologies) was added and the mixture incubated at 37°C for 10 minutes to allow ribonucleoprotein formation. These solutions for both genes were combined in a 1:1 ratio and 2-5 nL was injected in zebrafish embryos at the one-cell stage. Injected embryos were incubated in 20 mL of E3 embryo medium at 28°C until 24 hours post fertilisation (hpf).

The mutagenesis of *naglu* was performed similarly to as described by Fernandez et al. (2018). The sgRNAs were diluted to 24 μM in 1X TE buffer and mixed in a 1:1 ratio (1 μL:1 μL) with Cpf1 (Cas12a) nuclease (Integrated DNA Technologies) which had been diluted to 20 μM in Cas9 Working Buffer (20 mM HEPES, 300 mM KCl). This mixture was incubated at 37°C for 10 minutes, then ∼2-5 nL was injected into zebrafish embryos at the one-cell stage. Injected embryos were incubated in 20 mL of E3 embryo medium at 34°C until 24 hpf, before being moved back to 28°C.

#### Screening

At 24 hpf, 10 embryos from each injected group were removed for detection of mutagenesis. The embryos were incubated in 50 μL of recombinant Proteinase K (Roche Holding AG, Basel, Switzerland) diluted to ∼2 mg/mL in 1 x tris-EDTA (TE) at 55 °C for 3 hours, with each sample being agitated 3x during this period to disrupt the tissue. Proteinase K activity was inhibited by incubation at 95°C for 5 minutes before insoluble cellular debris was sedimented for 3 minutes at 16,000 g. The remaining genomic DNA (gDNA) was used for detection of mutation after PCR (see method below) using the primers described in **Table S2**.

PCR amplicons were tested for the presence of mutations using a T7 endonuclease I assay (New England Biolabs®, Ipswich, USA). For each PCR, 10 μL was subjected to electrophoresis on a 1% agarose gel according to the protocol below. This was done to verify that non-specific fragments from PCR amplification smaller than the target amplicon (and which could interfere in the interpretation of T7 cleavage products) were not present. Following this check, samples were split into two separate reaction tubes containing 8 μL of PCR product + 1 μL 10X NEB® Buffer #2 (New England Biolabs®). These were incubated in a thermocycler with an initial denaturation of 95 °C for 5 minutes. Heteroduplexes were then formed via annealing by lowering the temperature from 95 down to 85 °C at a ramp rate of -2 °C/second, followed by a ramp rate of -0.1 °C/second from 85 °C down to 25 °C. 0.5 μL of T7 Endonuclease I (New England Biolabs®) was then mixed into one tube of each sample. All tubes were incubated at 37 °C for 20 minutes to allow cleavage to occur, before reactions were halted by cooling the tubes to 4 °C. Samples were mixed with 0.2 volumes of 6X gel loading buffer (New England Biolabs®) and then electrophoresed in pairs alongside their undigested counterparts on an agarose gel as described below. An example gel image of the T7 assay for each mutant is shown in **Fig.S2**.

#### Sanger sequencing

Tail genomic DNA from fish determined to be homozygous (using the methods described in the **Genotyping** section below) for each of the three mutations was collected and subjected to PCR amplification as described below using primers flanking the mutation target site (**Table S2**). PCR reactions were resolved (see **Gel electrophoresis** section below) before the bands corresponding to the homozygous amplicon size were excised and purified using the Wizard® SV Gel and PCR Clean-Up System (Promega, Madison, Wisconsin, USA) according to the manufacturer’s protocol. DNA concentrations and the general quality of eluted samples were estimated on a NanoDrop® spectrophotometer (Thermo Fisher Scientific, Waltham, USA). Samples were prepared according to the recommended protocol of the Australian Genome Research Facility (AGRF, Adelaide, SA, Australia) before delivery to the AGRF. A single genotyping primer (**Table S2**) was used per sample according to which primer of a genotyping pair was more optimally located for sequencing relative to the mutation site; the Forward Primer for *sgsh*, the Forward Primer for *naglu* and the Reverse Primer for *hgsnat*. Samples were sequenced using Big Dye Terminator (BDT) chemistry version 3.1 (Applied Biosystems, Thermo Fisher Scientific).

#### *In silico* protein structure modelling

In the absence of experimentally derived crystal structures, the predicted wild type zebrafish protein structures were obtained from the AlphaFold database (Jumper et al., 2021; Varadi et al., 2022) and were compared to experimentally obtained human structures to identify important regions such as active sites and dimerisation interfaces (SGSH: PDB 4MHX, NAGLU: PDB 4XWH, HGSNAT: PDB 8TU9) (Birrane et al., 2019; Navratna et al., 2024; Sidhu et al., 2014). Mutant structures were predicted for zebrafish Sgsh^S387Lfs^ (ENSDART00000063147.5), Naglu^A603Efs^ (ENSDART00000048046.5), and Hgsnat^G577Sfs^ (ENSDART00000109911.5) using RoseTTAFold (Baek et al., 2021). Amino acid sequences were generated based on Sanger sequencing data (see above) and it was assumed that the deletion in *hgsnat* resulted in a continued read-through into the following intron, as the splice site was included in the deletion. These models were visualised aligned to the wild type protein structures using PyMOL (Schrodinger, 2015) (The PyMOL Molecular Graphics System, Version 3.0.0 Open-source, Schrödinger, LLC).

### Genotyping

Zebrafish were anaesthetised in 4 mg/mL tricaine methane sulfonate (Sigma-Aldrich®) before a 1 mm^2^ piece of tail fin was sliced off with a sterile scalpel. Tissue was incubated with 100 μL of recombinant Proteinase K (Roche) diluted to ∼2 mg/mL in 1 x tris-EDTA (TE) at 55 °C for 3 hours to release gDNA, with each sample being agitated 3x during this period to disrupt the tissue. Enzymatic activity was terminated at 95°C for 5 minutes before insoluble cellular debris was sedimented by centrifugation for 3 minutes at 16,000 g. Supernatants were stored at -20°C before use directly as template inputs for PCRs (see method below). PCR product(s) from each fish were resolved using gel electrophoresis (**Fig.S3**) to determine the genotype according to the method described below.

### Polymerase chain reactions

PCRs were performed using primers flanking the mutation sites of *sgsh, naglu* and *hgsnat*. These reactions were assembled to a final volume of 20 μL using 5X Green GoTaq® Reaction Buffer and dNTPs (Promega) with laboratory-purified DNA Taq Polymerase. PCR conditions consisted of an initial 2 minute denaturation at 95°C, followed by 35 cycles of: 30 seconds denaturation at 95°C and then 30 seconds annealing and extension at 72°C. Reactions were finalised with a 5 minute extension period at 72°C before storage at -20°C for later gel electrophoresis. Primer sequences, annealing temperatures, and extension times can be found in **Table.S2**.

### Gel electrophoresis

PCR products were loaded directly into ∼2.5-3% w/v agarose gels in 1X Tris-acetate-EDTA (TAE) buffer (for Sanger sequencing and T7 assays) or 1X sodium borate (SB) buffer (for genotyping) with 1.5 mg/mL ethidium bromide (Sigma-Aldrich®). Samples were run alongside 5 μL of 1kb Plus DNA Ladder (New England Biolabs®) or pHAPE ladder (Allen et al., 2023) and separated at 90V (TAE) or 200V (SB) for 30-90 minutes before visualisation under UV light. All our MPS III zebrafish mutations are deletions, so mutation-site-spanning PCR amplicons from wild type fish will appear after electrophoresis as a single higher molecular weight band, while amplicons from homozygous fish will appear as a single lower molecular weight band, and amplifications from heterozygous fish will display both bands. An example of our PCR genotyping assay is found in **Fig.S3**.

### Measurement of heparan sulfate (HS) in MPS III adult zebrafish brains

Adult zebrafish (3 months-of-age) were humanely euthanised in a loose ice slurry. Heads were removed by cutting at the level of the gills using sterile blades. The heads were then placed in petri dishes containing ice-cold 1 x PBS. Brains were carefully dissected from skulls and then placed, individually, into 500 µL aliquots of 10% methanol. The brains were then immediately homogenised on ice using a handheld pestle for two minutes. The brain homogenates were then sonicated using a Bioruptor^⍰^ (Diagenode, Denville, New Jersey, USA) bath sonicator for 10 minutes on high setting with cycles of 30 seconds on, 30 seconds off. After sonication, the brain homogenates were cleared by centrifugation at 13,000 g for 5 minutes at 4°C. Total protein concentrations in the supernatants were measured using the EZQ® Protein Quantitation Kit (Molecular Probes, Inc. Eugene, Oregon, USA) following the manufacturer’s protocol. Ten µg of total protein from each brain was delivered to the Mass Spectrometry Core Facility of the South Australian Health and Medical Research Institute (SAHMRI, Adelaide, SA, Australia) for measurement of disaccharide products of the butanolysis of HS (which is directly proportional to total HS) as described previously (He et al., 2019).

### Measurement of enzyme activity in MPS III adult zebrafish brains

To determine the level of remaining enzyme function in each MPS III model, fluorometric enzyme activity assays were performed on sibling fish resulting from heterozygous in-crosses of *sgsh*^*S387Lfs*^, *naglu*^*A603Efs*^ and *hgsnat*^*G577Sfs*^ respectively. Five individuals of each genotype (wild type, heterozygous and homozygous) were used from each mutant family. These adult zebrafish (aged approximately 12 months for *sgsh*, 24 months for *naglu* and *hgsnat*) were humanely euthanised in a loose ice slurry. Heads were removed by cutting at the level of the gills using sterile blades. These were then placed in petri dishes containing ice-cold 1 x PBS. Whole brains were carefully dissected from skulls and then snap frozen on dry ice. Homogenates were prepared in lysing Matrix D tubes (MP Biomedicals #116913500) filled with 0.5 mL of ice-cold 0.1% (v/v) TritonX-100 using a Pathtech Bead Bug tissue homogeniser. Enzyme activities were determined using fluorogenic substrates with minor modifications to the original descriptions of the methods (Karpova et al., 1996; Marsh and Fensom, 1985; Voznyi Ya et al., 1993). The total protein content was measured using MicroBCA Protein Assay kits (ThermoFisher Scientific #23225).

For Sgsh activity assays (Karpova et al., 1996; Whyte et al., 2015), 10 µL of sample was mixed with 20 µL of 5 mM 4-methylumbelliferyl *N*-sulpho-α-D-glucosaminide substrate dissolved in Michaelis’ barbital sodium acetate buffer, pH 6.5 (Carbosynth EM06602) and incubated at 47°C for 16 hours. After terminating the first reaction with 6 μL of twice-concentrated McIlvaine’s phosphate/citrate buffer, pH 6.7, 0.02% (w/v) sodium azide, the coupled reaction was conducted at 37°C for 24 hours using α-glucosidase (0.1 U in 10 μL; Sigma-Aldrich #G3651). The reaction was stopped on addition of 100 μL of stop buffer (0.5M Na_2_CO_3_/NaHCO_3_, pH 10.7).

For Naglu assays (Marsh and Fensom, 1985), 12.5 µL of sample was mixed with 12.5 µL of 2 mM 4-methylumbelliferyl-2-acetamido-2-deoxy-α-d-glucopyranoside substrate dissolved in 0.2 mM sodium acetate, pH4.3 (Carbosynth EM182013). Samples were incubated at 37°C for 16 hours and the reactions terminated on addition of 375 μL of stop buffer.

For Hgsnat activity (Voznyi Ya et al., 1993), 10 µL of sample was mixed with 10 µL of 6 mM acetyl coenzyme A trisodium salt and 10 µL of 3 mM 4-Methylumbelliferyl ⍰-D-glucosaminide dissolved in McIlvaine’s phosphate/citrate buffer, pH 5.6. Samples were also prepared with 10 µL of water in place of acetyl coenzyme A. Incubations were conducted at 37°C for 16 hours before adding 200 μL of stop buffer.

All enzyme reactions were clarified by centrifugation and supernatants were transferred to clear 96-well plates (Interpath #655101) which also included a 4-methylumbelliferone standard curve (Sigma-Aldrich #M1381). Fluorescence was measured at excitation 355 nm and emission 460 nm using a Spectramax iD5 plate reader. Data were expressed as the enzyme activities after correction of the substrate-blank counts and normalised for protein content. These data were subject to Welch’s One-Way ANOVA to account for unequal variance between groups, followed by Games-Howell *post-hoc* pairwise comparisons.

### Larval behaviour data collection

Two adult zebrafish heterozygous for MPS III mutations were in-crossed and the resulting progeny collected and housed together within petri dishes containing 50 mL of E3 medium (Westerfield, 2000) at 28.5 °C. Lighting conditions were set on a cycle of 14 hours in the light and 10 hours in the dark to simulate natural conditions. At 2 dpf, the larvae were transferred into a ∼ 26 x 16 cm tank and resuspended in a fresh batch of E3 media at a depth of 3 cm. At 4 dpf, larvae were placed individually into wells of clear Falcon 24-well plates (Corning, New York, United States) in 1 mL of E3 medium (**Fig.S4**).

The movement of larvae in each plate was recorded within the DanioVision Observation Chamber (Noldus, Wageningen, Netherlands) with the temperature control unit set at 28.5°C. The larvae within each plate were allowed to habituate to the behaviour room 30 minutes prior to testing. The movement of these experimentally naïve larvae was recorded using the DanioVision camera at 15 frames per second and 1280×960 resolution for one hour. Behavioural tracking was performed with EthoVision XT Version 11.5.1026 (Noldus). A 11.5 mm diameter circle was added inside individual wells to represent the central exploratory zone. The time spent moving was set at a starting threshold at 2 mm/s and stopping threshold at 1 mm/s and was averaged over 13 frames for all tests. Maximum velocity was set at 16 mm/s and samples were filtered out if exceeded. Videos were manually inspected to ensure proper tracking. Then, the overall distance travelled, distance per specified time bin, time and crossings into the central zone data was exported into a spreadsheet which contained these parameters summed over three 20 minute time bins. After behavioural data was collected, individual larvae were euthanised in ice water and genomic DNA extracted. Samples were then genotyped using PCRs and gel electrophoresis.

### Analysis of larval behaviour data

All statistical analysis was performed in *R* (R Core Team, 2021). To determine whether MPS III genotype had a significant effect on the total distance travelled (a marker of activity), the time spent in the centre and the frequency of crossings into the centre zone, we fitted these data to linear mixed effect models using *lme4* (Bates et al., 2015), logit-linked beta generalised linear models with *glmmTMB* (Brooks et al., 2017), and log-linked negative binomial models respectively.

In all models, we specified genotype, time bin and an interaction of genotype and time bin as fixed effects, and behaviour trial and a fish identifier (to account for repeated measures) as random effects. In the total distance travelled dataset, model assumptions were violated for all MPS III subtypes. Subsequently, the distance travelled response variables were transformed by square root or log10 so that model assumptions were no longer violated. The time spent in the centre and time spent moving data were transformed into ratios of each time bin by dividing by 1200. All zero values were converted to an extremely small positive value before being fitted to logit linked beta models. The significance of the fixed effects in all models were determined with a Wald type II *χ*^2^ test (implemented in the *car* (Fox and Weisberg, 2019) package) with a threshold of significance at p-value < 0.05. Estimated marginal means were calculated using *emmeans* (Russell V. Lenth, 2023).

### Zantiks Free Movement Pattern Y-maze

#### Data collection

We created families of siblings with wild type, heterozygous, and homozygous genotypes simultaneously by in-crossing pairs of fish heterozygous for each mutation (**Fig.4A**). Due to some difficulties with embryo collection, multiple mating events per parent pair were required to produce the 100 individual fish required for testing. Families possessing the same mutation were raised side-by-side in tanks within the same recirculated water aquarium system.

**Fig 1:**
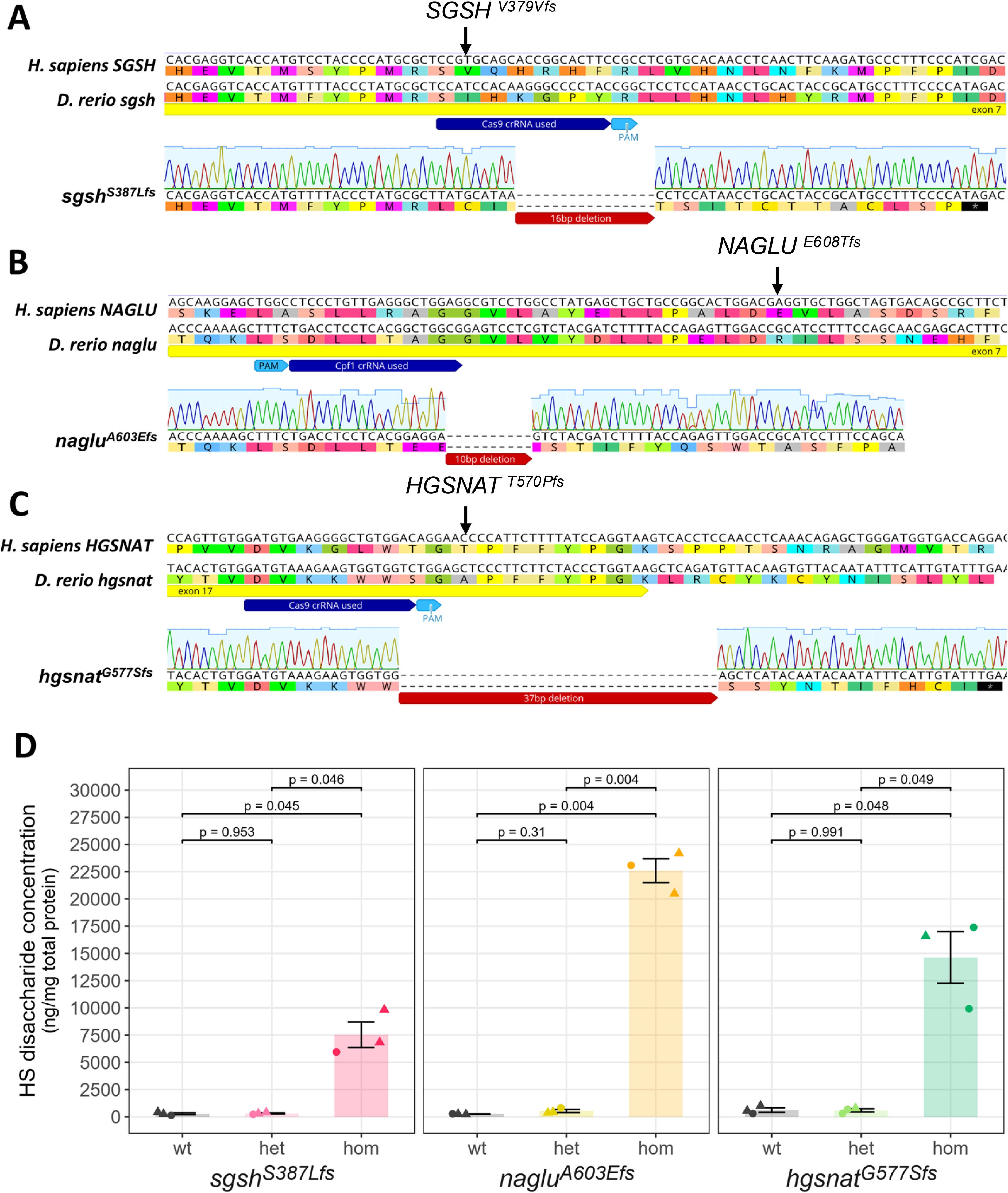
Summary of MPS III mutant characterisation. **A-C**, Sanger sequencing chromatograms of PCR amplicons from homozygous mutant zebrafish (*D. rerio*). **A, *sgsh***^***S387Lfs***^**; *B, naglu***^***A603Efs***^; and **C, h*gsnat***^***G577Sfs***^ zebrafish sequences, aligned to the corresponding wild type genomic sequences of *Homo sapiens* and zebrafish. Light blue shading on each chromatogram indicates the sequencing base call quality. CRISPR guide RNAs (dark blue), PAMs (light blue), exonic regions (yellow), and deletion sites (red) are indicated. Amino acid sequences are displayed under each nucleotide sequence. Known human mutation sites are annotated. **D**, Mass spectrometric measurement of heparan sulfate (HS) disaccharide concentration in wild type, heterozygous and homozygous 3-month old *sgsh*^*S387Lfs*^ (left panel), *naglu*^*A603Efs*^ (centre panel) and *hgsnat*^*G577Sfs*^ (right panel) zebrafish brains. Data is presented as mean ± SEM and tested via Welch’s ANOVA (**Table S3**) and Games-Howell pairwise comparisons (shown on figure). Data points represent a single measurement from an individual brain (triangular for males, circular for females). Raw mass spectrometric data can be found in **Supplemental Data File 1**.

**Fig 2:**
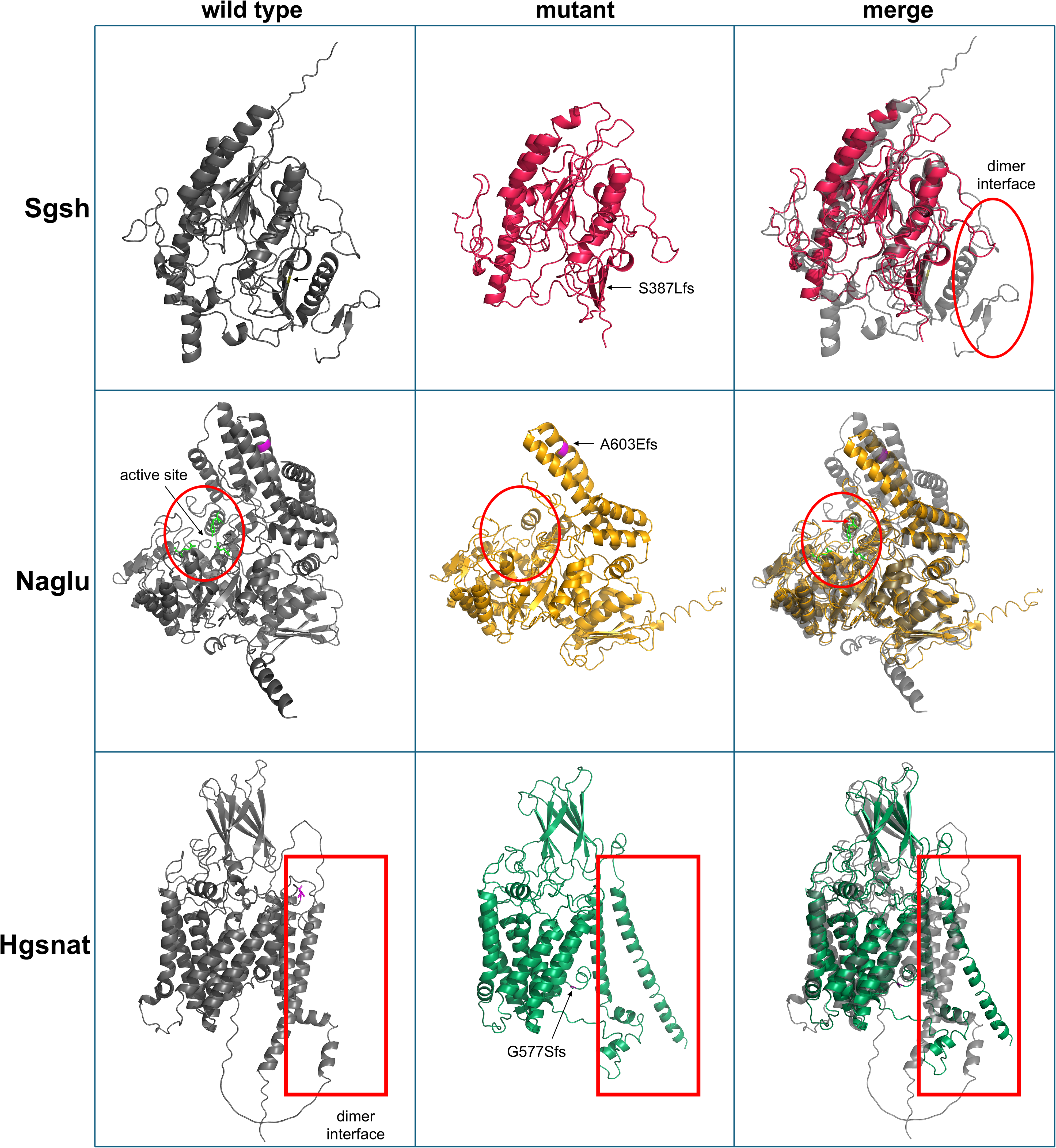
*in silico* modelling of MPS III zebrafish protein structures. AlphaFold-predicted wild type (left column) and RoseTTAFold-predicted mutant (centre column) protein structures for Sgsh^S387Lfs^ (top row), Naglu^A603Efs^ (centre row), and Hgsnat^G577Sfs^ (bottom row). PyMOL-aligned structures are shown in the right column. The locations of each mutation are indicated by black arrows and highlighted in yellow (Sgsh)(obscured from this viewpoint) and pink (Naglu, Hgsnat). Features of interest include dimerisation interfaces for Sgsh and Hgsnat (enclosed in red) and stabilising residues around the active site in Naglu (highlighted green), appearing to be affected in the mutant (red arrow, merged image). The per-residue estimated error for each of these structures is presented in **Fig.S14**.

**Fig 3:**
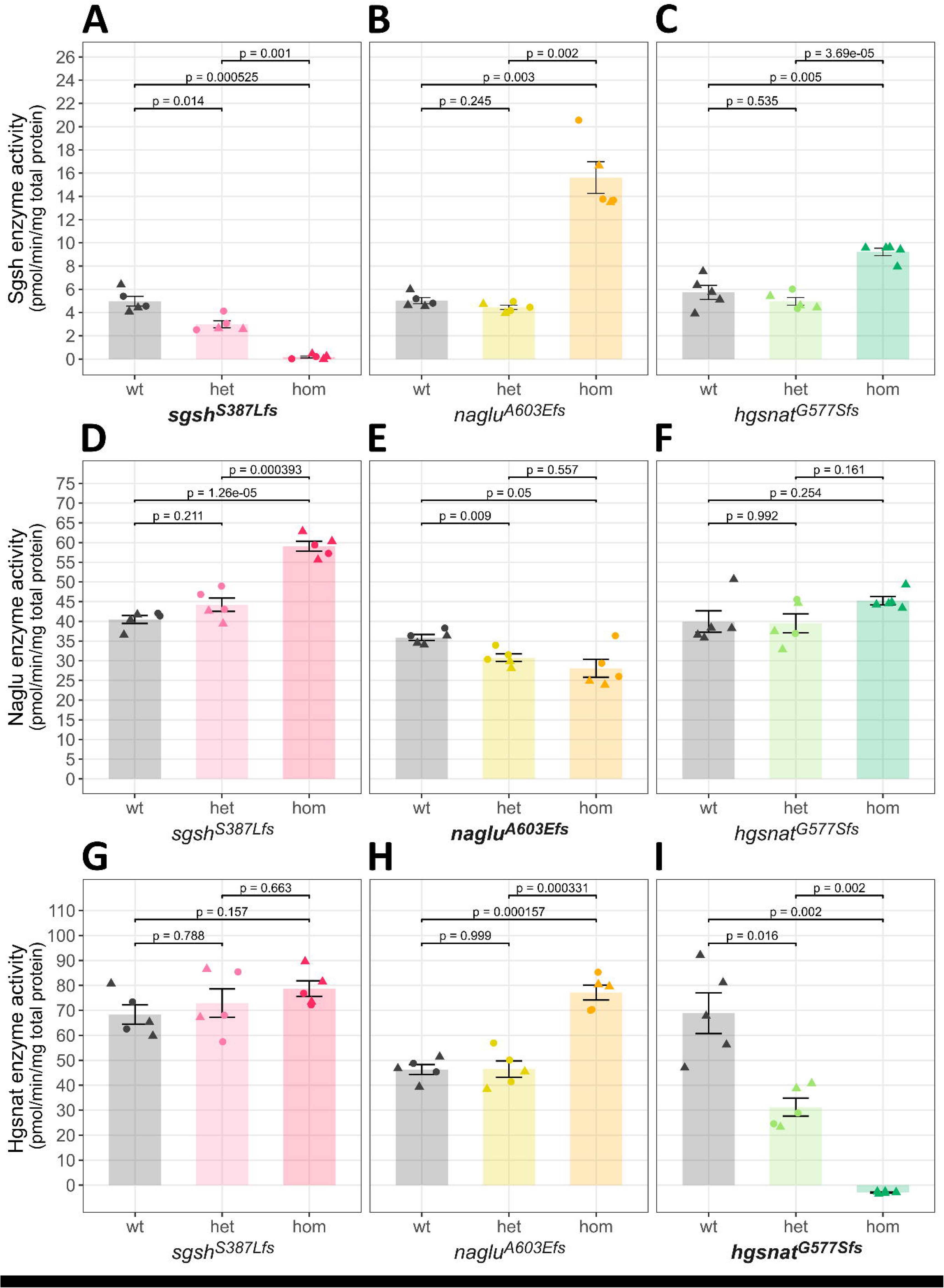
MPS III zebrafish mutations significantly affect enzyme activity levels in fluorometric assays. **A-C**, Sgsh activity in *sgsh*^*S387Lfs*^ (**A**), *naglu*^*A603Efs*^ (**B**) and *hgsnat*^*G577Sfs*^ (**C**) families. **D-F**, Naglu activity in *sgsh*^*S387Lfs*^ (**D**), *naglu*^*A603Efs*^ (**E**) and *hgsnat*^*G577Sfs*^ (**F**) families. **G-I**, Hgsnat activity in *sgsh*^*S387Lfs*^ (**G**), *naglu*^*A603Efs*^ (**H**) and *hgsnat*^*G577Sfs*^ (**I**) families. Each panel shows enzyme activity in whole brain samples of wild type (wt), heterozygous (het) and homozygous (hom) fish from that respective family. Enzyme activity is plotted as pmol of substrate converted per min per mg of total protein. Data is presented as mean ± SEM and was tested via Welch’s ANOVA (**Table S3**) and Games-Howell pairwise comparisons. Data points represent a single measurement from an individual brain.

**Fig 4:**
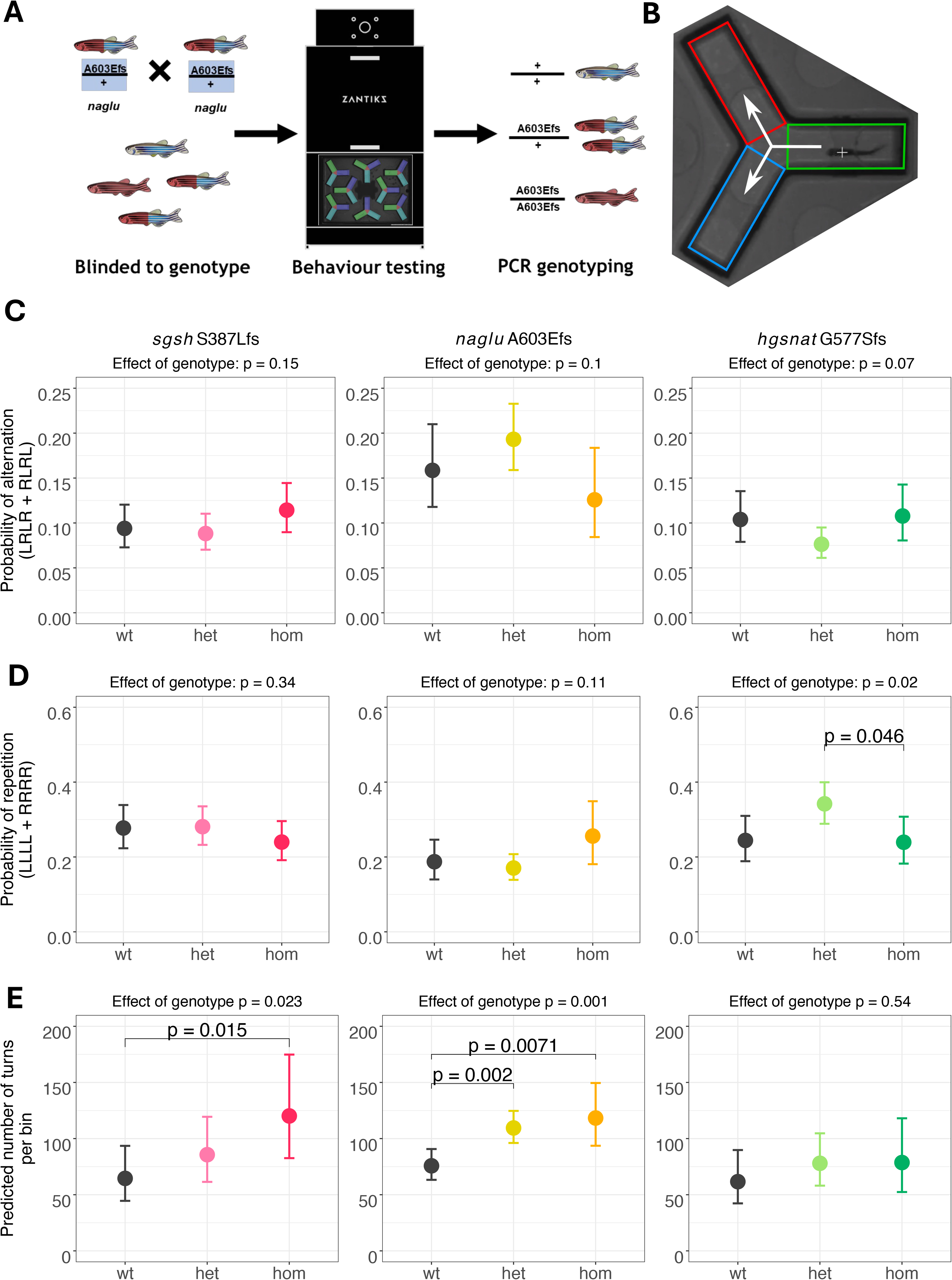
Adult MPS III zebrafish show hyperactivity in a Y-maze. **A**, The heterozygous in-cross breeding strategy used, by which experimental families are generated from a single parent pair. All genotypes of interest are present within each family, and the genotype of each individual fish is determined after behavioural testing is concluded. This figure displays this strategy as used to analyse the *naglu*^*A603Efs*^ mutation and was identical for analysis of the other mutations. **B**, Diagram of a single Y-maze arena divided into three zones. Left-right turn choices are recorded. **C–D**, Generalised linear mixed model–predicted probabilities of zebrafish performing an alternation (LRLR or RLRL; **C**) or a repetition (LLLL or RRRR; **D**) tetragram in the Y-maze. Fish were assessed at 20 months (*sgsh* S387Lfs; left panel), 21 months (*naglu* A603Efs; middle panel), and 17 months (*hgsnat* G577Sfs; right panel). Plotted values represent estimated marginal means. Values shown in the plot titles indicate the significance of the genotype effect (Type-II Wald χ^2^ test; all fixed effects listed in Table S3). Post-hoc Tukey contrasts are displayed only when *p* < 0.**0**5. **E**, Generalised linear model-predicted average number of turns per 10 minute time bin in the same behavioural experiments. Raw values can be found in **Fig.S17**.

We performed the Free-Movement-Pattern Y-maze using the Zantiks LT automated behavioural testing environment (Zantiks, Cambridge, UK) on our heterozygous in-cross families at 20 (*sgsh*^*S387Lfs*^), 21 (*naglu*^*A603Efs*^) and 17 (*hgsnat*^*G577Sfs*^) months of age. To reduce the confounding effects of gross circadian rhythm differences, we performed behavioural testing between 9am and 5pm, firmly within the zebrafish wake cycle. The testing of each family took place over the span of a single day. Following the protocol described previously (Cleal et al., 2021b), fish were isolated eight at a time for 30 minutes in the behavioural testing room to acclimatise to the new room environment. Then, they were placed in the Y-mazes containing 6 L of system water in the Y-maze-containing tank at an initial temperature of 27 °C, and allowed to swim freely in lit conditions (approximately 300-400 lux) for 1 hour in the absence of any external stimuli. Fish movement was tracked using the in-built Zantiks tracking system, which exports a video recording, and generates a spreadsheet containing the time points at which each fish enters and exits a zone of the Y maze.

#### Behavioural data analysis

All behavioural analysis was performed using the *R* programming language (R Core Team, 2021). Briefly, raw data files were processed using an *R* batch script (Fontana et al., 2019) and available from GitHub (https://github.com/clay-j/ZANTIKS_YMaze_Analysis_Script). The output of this script gives the frequencies of 16 tetragrams (four consecutive arm entries, consisting of all possible left (L) or right (R) turn choice combinations from LLLL to RRRR) over six time intervals (bins) of 10 minutes each. Data visualisation was performed using the *ggplot2* package (Wickham, 2016).

To assess any changes in the frequencies of the alternation tetragrams (LRLR and RLRL tetragrams) performed by our zebrafish in the Y-maze, we fitted the tetragram frequency data to a generalised linear mixed effects model using the *glmmTMB* package (Brooks et al., 2017) with a logit link function. We employed the beta-binomial distribution to account for overdispersion in the dataset. Random effects were a fish identifier number (e.g. fish 1, fish 2 etc.), and the start time of the behavioural test. Fixed effects were specified to be genotype (wild type, heterozygous, or homozygous), whether a fish displayed a behavioural lateralisation (left bias, right bias or no bias), sex, time bin during the Y-maze test, and an interaction effect between genotype and time bin. To test whether these fixed effects significantly impacted the alternation probability of the fish, we used Type II Wald *χ*^2^ tests on the generalised linear model using the *Anova* function of the *car* package (Fox and Weisberg, 2019). Fixed effects were considered to be significant if they had a p-value < 0.05. Estimated marginal means were calculated using *emmeans* (Russell V. Lenth, 2023) and *post-hoc* contrasts were performed using the Tukey method.

To assess whether our zebrafish models showed any changes in general activity or locomotion, we also calculated the total number of arm entries (turns) performed by each fish in the Y-maze over the six time bins. This data was fitted to another generalised linear model with a negative binomial distribution with a logit link function using the *glm*.*nb* function of the *MASS* package (Venables and Ripley, 2002). Random effects were fish ID and start time, while included fixed effects were genotype, time bin, clutch, sex and an interaction effect between genotype and time bin. As above, we used Type II Wald *χ*^2^ tests using the *Anova* function, with effects considered to be significant at a p-value < 0.05. Estimated marginal means were calculated as above.

### Transcriptome analysis

#### Data generation

To compare brain transcriptomes in our three MPS III zebrafish lines, we generated families of 2^nd^ Filial (F2) generation zebrafish originating from G_0_ in-crosses of parental zebrafish heterozygous for two of the MPS III mutations (**Fig.5A, Fig.S5-7**). We initially aimed to analyse n = 6 fish per genotype from single families of progeny from in-crosses of doubly heterozygous mutant fish. We chose n = 6 biological replicates per group, informed by our previous findings that this sample size yields ∼70-80% power even in models with subtle, early-onset transcriptomic changes (Barthelson, Karissa et al., 2020; Barthelson et al., 2025). Given that the biological effect sizes in the present study were anticipated to be considerably larger, this approach ensures high sensitivity for detecting differentially expressed genes. However, due to smaller than expected clutch sizes, we were forced to use multiple families of siblings, or n < 6, in some analyses. All RNA samples included in the final analyses met accepted quality thresholds for transcriptomic profiling, with the majority exhibiting RNA integrity numbers (RIN) >7. For the MPS IIIA and MPS IIIB cohort, we analysed n = 4 fish per genotype, which also included 4 *sgsh* ^*S387Lfs/+*^ fish to explore cellular processes affected in the brains of presumably unaffected carriers of MPS III mutations. For the MPS IIIA and MPS IIIC cohort, we sequenced all homozygous *sgsh* ^*S387Lfs*^ zebrafish in the family (n = 7) to account for the lower number of fish with this genotype in the MPS IIIA and MPS IIIC cohort. For the MPS IIIB and MPS IIIC cohort, we could not obtain a sufficiently large enough family in a single breeding event within our timeframe to be able to compare this with the other families. Therefore, we analysed fish spawned by two pairs of parents (P1 and P2, both doubly heterozygous for the *naglu*^*A603fs*^ and *hgsnat*^*G577Sfs*^ mutations) in a total of 3 spawning events. This information is summarised in **Fig.S5-7**.

**Fig 5:**
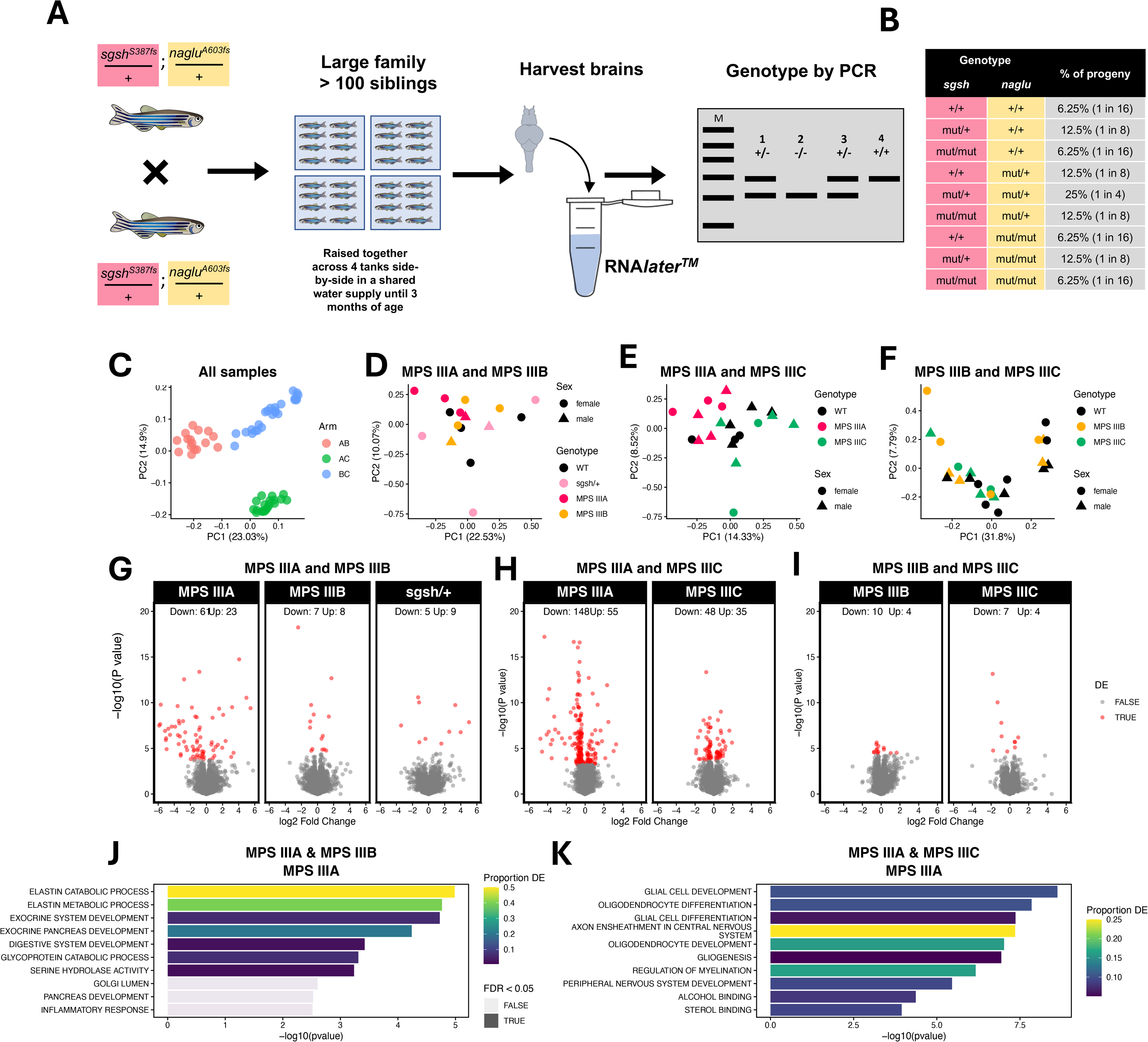
Intra-family brain transcriptome analysis. **A**, Zebrafish doubly heterozygous for two MPS III mutations (*sgsh* ^*S387Lfs*^ and *naglu* ^*A603fs*^ shown here) are in-crossed to generate a large family of at least 100 siblings. The family is raised across several tanks until 3 months of age. At that point, all fish are euthanised and their brains harvested and preserved in RNA*later* solution. Each fish is genotyped using PCR tests. **B**, Expected proportions of mutant genotypes assuming normal Mendelian inheritance (illustrated using *sgsh* and *naglu*). **C-F**, Principal component analysis of zebrafish brain transcriptome datasets at 3 months of age for **C**, all samples **D**, MPS IIIA and MPS IIIB cohort; **E**, MPS IIIA and MPS IIIC cohort; and **F**, MPS IIIB and MPS IIIC cohort. Each point represents a brain transcriptome, which is coloured according to genotype and shaped according to sex. The percentage of variance explained by principal components (PC) 1 and 2 are indicated within the x and y axis labels.**G-I**, Volcano plots of differential gene expression analyses. Plots are constrained between -6 and 6 on the x axis and between 0 and 20 on the y axis for visualisation purposes. Full outputs of differential expression testing can be found in **Supplemental Data File 2. J**, Top 10 gene ontology terms significantly over-represented in MPS IIIA zebrafish in the MPS IIIA and MPS IIIB cohort, and **K**, MPS IIIA and MPS IIIC cohort. The colour of the bars represent the proportion of the genes in the GO term which were differentially expressed in MPS IIIA zebrafish. Bars appear transparent if the FDR-adjusted p-value did not reach < 0.05. All significant GO terms in **I** can be found in **Fig.S18**.

Each family of fish was raised until 3 months of age, at which time the entire family was humanely euthanised in an ice slurry. Entire heads were excised at the level of the gills and were each incubated in 600 µl of RNA*later*™ Stabilization Solution (Invitrogen, Thermo Fisher Scientific, Waltham, USA) overnight at 4°C, before storage at -80°C until use. The tail of each fish was also removed for genomic DNA extraction and genotype determination by PCR as described above. Total RNAs were isolated from RNA*later-*preserved brains using the *mir*Vana™ miRNA Isolation Kit (Ambion, Life Technologies, Thermo Fisher Scientific, Waltham, USA) following the manufacturer’s protocol. To remove any carried-over genomic DNA during RNA purification, we treated the total RNA samples with *DNaseI* using the DNA-*free*™ DNA Removal Kit (Invitrogen, Thermo Fisher Scientific, Waltham, USA) following the manufacturers protocol for routine DNase treatment. Then, 500 ng of high-quality total RNA (with RIN_e_ generally ≥ 8) was delivered on dry ice to the South Australian Genomics Centre (SAGC, Adelaide, Australia) for preparation of stranded, polyA+ libraries using Nugen Universal Plus mRNA-seq (NuGEN, Ltd., UK). Cluster generation was performed using the Illumina to MGI Library Conversion (MGIEasy Universal Library Conversion Kit, Part No. MGI1000004155). Then, 2 x 98bp paired end sequencing was performed including 8 bp unique molecular identifiers (UMIs) using MGI DNBSEQ-G400 chemistry (MGI Tech, Shenzhen, China). Each of the libraries was sequenced over multiple lanes and the data from each library subsequently combined.

#### Pre-processing

Pre-processing was performed using a custom pipeline implemented in *snakemake* (Mölder et al., 2021). Briefly, the raw fastq files were subjected to initial quality checks using *fastQC* version 0.11.9 (Andrews, 2010). Then, read were filtered and trimmed using *fastp* version 0.23.1 (Chen et al., 2018). Reads containing more than 40% bases with Phred quality < 15 or more than 5 ambiguous bases (N) were removed. Adapter sequences were automatically detected and trimmed. Reads shorter than 15 bp after filtering were discarded. The remaining reads were aligned to the zebrafish genomes e (GRCz11, Ensembl release 101 (Cunningham et al., 2022)) using the splice-aware alignment software *STAR* version 2.7.0d (Dobin et al., 2013). Aligned reads associated with the same unique molecular identifier (UMI) correspond to PCR duplicates generated using library preparation. These were deduplicated using the *dedup* function of *umi_tools* v1.0.1 (Smith et al., 2017). A gene-level counts matrix was then generated using *featureCounts* version 2.0.1 (Liao et al., 2014).

#### Analysis

Statistical analysis of the RNA-seq data was performed using *R (R Core Team, 2021)*. Lowly expressed genes are considered uninformative for differential expression analysis. Therefore, for each experimental comparison, we established a counts per million threshold value that equated to 10 counts in the sample with smallest library size. Genes were removed if their expression was lower than this value in at least the number of samples equalling the smallest genotype group (as recommended in (Chen et al., 2016)). The remaining genes (i.e. above the detection threshold) were normalised using the trimmed mean of M-values (TMM) method (Robinson and Oshlack, 2010). Principal component analysis was performed to assess whether batch effects showed significant influence on brain transcriptomes. The factors of “Cohort” and “Home tank” were found to show effects on the brain transcriptome (**Fig.5C, S8-S10**). Therefore, each experimental cohort was analysed separately. “Home Tank” and “sex” was included in the design matrices for differential expression analyses. Differentially expressed (DE) genes were identified using generalised linear models and likelihood ratio tests in *edgeR* (Robinson et al., 2010). We considered a gene to be differentially expressed (DE) if the FDR-adjusted p-value was less than 0.05.

In initial differential gene expression analyses, we noted some small biases for genes to be called DE due to their %GC content and length (**Fig.S11-13**). Therefore, we used the conditional quantile normalisation (CQN, (Hansen et al., 2012)) method to account for this bias. For CQN, we used the average length (in base pairs) of transcripts per gene, and a weighted average (by transcript length) of the %GC content of detected transcripts per gene to generate an offset term which was included in DE analyses in *edgeR*.

For gene set analyses, we used the KEGG (Kanehisa and Goto, 2000) and gene ontology (GO, (Ashburner et al., 2000)) gene sets obtained from the Molecular Signatures Database (Subramanian et al., 2005) using *msigdbr* (Dolgalev, 2021). The KEGG and GO term gene sets were filtered to retain only genes detected in each cohort. The GO terms themselves were also filtered so that only GO terms with more than 3 steps to the root node were retained to reduce redundancy. The KEGG gene sets were additionally filtered to retain only gene sets containing more than 5 genes. To test for changes to iron homeostasis, we used the lists of genes identified to contain iron-responsive elements (IREs) in the untranslated regions of their encoded transcripts from (Hin et al., 2021). To test for changes in cell type proportions, we used the marker genes of adult zebrafish brain cell types (23 different cell types) from a single-cell RNA-seq dataset (Jiang et al., 2021). Marker gene sets were defined by whether the gene showed an average logFC in the cell type of interest greater than 1.5 relative to all other cell types in the Jiang et al. dataset.

We tested whether the GO gene sets were enriched with DE genes using *goseq* (Young et al., 2010) specifying the co-variate as the average transcript length per gene. We considered a GO term to be significantly enriched if it contained at least 2 DE genes, and the FDR-adjusted p-value was less than 0.05. We also utilised the fast implementation of the *ROAST* algorithm: *fry* (Wu et al., 2010) for the KEGG, cell type marker and IRE gene sets. We considered a gene set to be significantly altered as a group if the FDR-adjusted p-value was < 0.05.

*Post-hoc* power analysis was performed using the *ssizeRNA_vary*() function of the *ssizeRNA* package (Bi and Liu, 2016). The calculations were performed at the controlled false discovery rate of 0.05, with a pseudo sample size of 40 and under the assumption that 90% of the genes were not differentially expressed. The mean gene counts and dispersions in the control group were set from those calculated in each individual cohort after adjustment for CQN.

### Data and code availability statement

The RNA-seq datasets generated in this study has been deposited to the GEO database (accession number GSE244310). All code to reproduce the transcriptome analysis can be found at https://github.com/karissa-b/2022_MPSIII_3mBrainRNAseq. Larval behaviour analysis can be found at https://github.com/Lakshay-Rattan/Honours_MPSIII_Analysis. Adult behaviour analysis can be found at https://github.com/ewan-gerken/MPSIII_zebrafish_Ymaze_aged. Biochemistry analyses can be found at https://github.com/ewan-gerken/MPSIII_zebrafish_enzymeassays and https://github.com/ewan-gerken/HS_accumulation_MPSIII_3m.

## Results

### Creation of zebrafish models of MPS IIIA, B and C

Disease mutations disrupting translational reading frames are known to exist in the most downstream exons of the human *SGSH* (Yogalingam and Hopwood, 2001), *NAGLU* (Yogalingam and Hopwood, 2001), and *HGSNAT* (Feldhammer et al., 2009) genes. We sought to generate similarly disruptive frameshift mutations in the respective orthologous zebrafish genes *sgsh, naglu* and *hgsnat*, which exhibit significant conservation with their human counterparts. Specifically, Sgsh shares 69.4% identity (87.1% BLOSUM45 similarity) with the human protein, while Naglu and Hgsnat share 56.5% identity (84.5% similarity) and 54.1% identity (77% BLOSUM45 similarity), respectively. The advantage of identifying frameshift mutations leading to stop codons in the final exons of these genes is that such mutations are less likely to cause nonsense-mediated decay (NMD) of mutant mRNAs and so may avoid transcriptional adaptation (El-Brolosy et al., 2019) that can complicate the interpretation of mutant phenotypes. Using CRISPR-Cas and CRISPR-Cpf1 systems, we introduced the mutations *sgsh*^*S387Lfs*^, *naglu*^*A603Efs*^, and *hgsnat*^*G577Sfs*^ (**Fig.1A-C**). These mutations show similarity to the human mutations *SGSH*^*V37V9fs*^ (Muschol et al., 2004), *NAGLU*^*E608Tfs*^ (no published cases, predicted to be pathogenic in ClinVar, accession RCV002007460.5), and *HGSNAT*^*T570Pfs*^ (observed as one *HGSNAT* allele in a case of adult-onset nonsyndromic retinitis pigmentosa, (Schiff et al., 2020)). Importantly, all of our zebrafish mutations cause frameshifts upstream of the equivalent positions of known rapidly progressing human MPS III mutations (**Fig.S1**), and so presumably are similarly detrimental to the enzyme activities of the genes’ protein products.

We next aimed to estimate what effect these mutations may have on the three-dimensional structure of their respective encoded proteins. To achieve this, we performed *in silico* modelling using RoseTTAFold (Baek et al., 2021) to predict the structure of each mutant protein and align them with the wild type structure. structure. These comparisons revealed considerable changes to the proteins’ tertiary structures (**Fig.2**). In Sgsh and Hgsnat, the truncations resulted in significant alterations to their dimerisation interfaces, potentially reducing their oligomeric stability and ability to degrade HS. The changes in the Naglu structure appeared to affect the active site with truncation of the protein resulting in the loss of stabilising residues which is predicted to also reduce this protein’s ability to degrade HS.

### Zebrafish models of MPS IIIA, B and C show biochemical characteristics consistent with the human disease

HS accumulation is the primary pathological hallmark of MPS III. Therefore, we first measured the butanolysis disaccharide product of HS (directly proportional to HS (Trim et al., 2015)) in the brains of 3-month-old zebrafish by mass spectrometry (He et al., 2019). In all three homozygous mutants, HS levels were significantly elevated (**Fig.1D**), measuring 26-fold higher for *sgsh*^*S387Lfs*^ (p = 0.046), 85-fold higher for *naglu*^*A603Efs*^ (p = 0.004) and 23-fold higher for *hgsnat*^*G577Sfs*^ (p = 0.048). Zebrafish brains heterozygous for each mutation showed HS levels unchanged from basal wild type levels, consistent with the observation that carriers of loss-of-function alleles in lysosomal enzymes do not develop lysosomal storage disorders (Douglass et al., 2021). Together these data show that all three mutants recapture the primary MPS III hallmark and indicate that the activities of the enzymes responsible for HS breakdown are meaningfully impacted.

We next aimed to confirm to what degree our MPS III mutations impacted the enzyme function encoded by their respective genes using fluorometric assays. Sgsh enzyme activity was reduced in *sgsh*^*S387Lfs*^ homozygous fish, to a mean of 3.6% of wild type levels (p = 0.0005). Sgsh activity in heterozygous fish within this family was also significantly different compared to both wild type and homozygous fish, consistent with a typical genetic dose response (**Fig.3A**). In *hgsnat*^*G577Sfs*^ homozygous fish, we observed a complete loss of Hgsnat activity to below the limit of detection in these samples (p = 0.002, **Fig.3I**), with a reduction of approximately 50% in heterozygous samples akin to that observed in heterozygous *sgsh*^*S387Lfs*^. Upon measuring Naglu enzyme activity, we observed comparatively modest, yet statistically significant, reductions in both heterozygous and homozygous samples. Mean activity was measured at approximately 78% of wild type levels (p = 0.05, **Fig.3E**) in *naglu*^*A603Efs*^ homozygous brains.

The lysosomal enzyme activity profiles for Sgsh, Naglu and Hgsnat were also examined in fish carrying mutations in the genes not directly associated with those enzyme activities. Sgsh activity was significantly elevated in fish homozygous for *naglu*^*A603Efs*^ (**Fig.3B**) and *hgsnat*^*G577Sfs*^ (**Fig.3C**), with 3.1 and 1.6-fold increases respectively, relative to wild type levels (p = 0.003 and p = 0.005). A similar 1.5-fold increase from wild type levels was found for Naglu activity in *sgsh*^*S387Lfs*^ homozygous fish (p = 1.26E-05, **Fig.3D**), with a slight yet insignificant increase in mean Naglu activity for homozygous *hgsnat*^*G577Sfs*^ (**Fig.3F**). Mean Hgsnat activity in *sgsh*^*S387Lfs*^ homozygous fish also trended upwards (**Fig.3G**), however this was again insignificant. Finally, *naglu*^*A603Efs*^ homozygous brains showed a 1.7-fold increase in Hgsnat activity relative to wild type (p = 1.6E-04, **Fig.3H**).

### Zebrafish models of MPS IIIA and B show hyperactivity in adulthood

Children with MPS III show severe hyperactivity early in disease progression (Meyer et al., 2007). To determine whether the MPS III zebrafish larvae also show this disease characteristic, we assessed their swimming behaviour in an open field at 5 dpf. Overall, zebrafish of each genotype moved less with increasing time (i.e., increasing bins), consistent with habituative behaviour. However, we did not observe any statistically significant effect of genotype in any MPS III model on their distance travelled, or their time spent moving (**Fig.S15, 16**). Therefore, MPS III larvae do not appear to exhibit hyperactivity at 5 dpf.

The MPS IIIA zebrafish model of Douek et al. (2021) was found to exhibit less thigmotactic behaviour at 5 dpf, interpreted as reduced anxiety (Valle, 1970). Here, only homozygous *sgsh*^*S387Lfs*^ larval zebrafish exhibited increased thigmotactic behaviour, particularly in the first 40 minutes of the test (**Fig.S15, 16**). Another measurement of thigmotactic behaviour is the number of crossings into the central zone. Modelled data demonstrated no significant effect of genotype (**Fig.S15, 16**). Therefore, it is unlikely that robust differences in anxiety-related behaviour emerge by 5 dpf in any of the three MPS III models.

We next evaluated learning and memory in adult MPS III zebrafish using the Free Movement Pattern (FMP) Y-maze assay (Cleal et al., 2021b), a validated measure of short-term spatial working memory used in several zebrafish studies (Cleal et al., 2021a; Cleal et al., 2021b; Fontana et al., 2019; Fontana et al., 2020; Fontana et al., 2021). During a one hour trial, sequential left and right turn choices are recorded (**Fig.4A, B**) and parsed into overlapping four-turn sequences (tetragrams), with alternation patterns (LRLR and RLRL) representing the preferred search strategy and serving as the primary readout of working memory. The sensitivity of this assay to working memory has been established previously, as pharmacological disruption of neurotransmitters important for memory (glutamate, acetylcholine, and dopamine) abolishes alternation preference behaviour (Cleal et al., 2021b). Repetitive tetragrams (LLLL and RRRR) are interpreted as behavioural perseveration, reflecting a reduced tendency to shift response strategies and a bias toward rigid turning behaviour, consistent with impaired cognitive flexibility (Fontana et al., 2019). Overall activity between groups can also be assessed by comparison of the total number of turns performed.

We assessed the spatial working memory (alternations), behavioural perseveration (repetitions) and overall activity (total turns) of our MPS III zebrafish at using the FMP Y-maze test. Each zebrafish family, aged approximately 20 months, was composed of wild type, heterozygous, and homozygous siblings (**Fig.4A, Fig.S17**). Across all MPS III models, we found no significant effects of genotype on the estimated probability of fish performing alternating turn choices alternation (**Fig.4C**). Therefore, there is no evidence supporting altered spatial working memory in these three MPS III mutants at the age tested. A significant effect of genotype on repetitions was observed within the *hgsnat* cohort. However, *post-hoc* comparisons revealed that this difference was driven by the contrast between heterozygous and homozygous fish (**Fig.4D**). The *hgsnat*^*G577Sfs*^ homozygous mutants exhibited levels comparable to wild-type controls, suggesting that this effect does not represent a disease-associated phenotype.

To assess whether our three zebrafish MPS III models might show the hyperactivity characteristic of the human condition, we examined the number of turns made during one hour in the Y-maze. We found that the effect of genotype on the number of turns predicted by the model was significant for *sgsh* and *naglu* mutant cohorts (**Fig.4E, Table S4**). *Post-hoc* comparisons revealed a significant hyperactivity phenotype in homozygous *sgsh*^*S387Lfs*^ (p = 0.015) and homozygous *naglu*^*A603Efs*^ (p = 0.0071) zebrafish relative to their wild type siblings. Unexpectedly, heterozygous *naglu*^*A603Efs*^ mutant zebrafish also showed a significantly increased activity (p = 0.002), suggesting that this mutant genotype may also have a hyperactivity phenotype. The *hgsnat*^*G577Sfs*^ mutant zebrafish did not show statistically significant changes in activity during this experiment.

### Brain transcriptome analysis of zebrafish models of MPS IIIA, IIIB and IIIC reveals common dysregulated KEGG pathways

One motivation behind construction of our zebrafish models of MPS IIIA, MPS IIIB, and MPS IIIC mutations was to characterise the molecular state of the MPS III brain. Transcriptome analysis is currently the single technique giving the greatest volume of molecular phenotypic data from analysis of cells and tissues. Therefore, we sought to apply transcriptome analysis to characterise the molecular state of brains in the three models. We enhanced the sensitivity of this analysis to detect differences between MPS III types by applying our intra-family strategy of genetic and environmental noise reduction (**Fig.5A,B**). The intra-family analysis strategy also facilitated an experimental structure incorporating internal replication; we bred families allowing comparisons of MPS IIIA and MPS IIIB to wild type, MPS IIIA and MPS IIIC to wild type, and MPS IIIB and MPS IIIC to wild type. We analysed brains at 3 months of age, a developmental stage at which zebrafish are approaching sexual maturity (Parichy et al., 2009), the brain is structurally mature (reviewed in (Vaz et al., 2019)), and approximately equivalent to late childhood in humans (McElroy et al., 2016). Additionally, we had already detected hyperactive behaviour at a later age and our previous transcriptome analyses of familial Alzheimer’s disease-like mutations (expected to have much milder phenotypes than those causing MPS III) at 6 months of age had detected significant differences in gene expression (Barthelson et al., 2022).

We first performed principal component analysis (PCA) on the RNA-seq data to assess the overall similarity between samples. In the entire dataset, samples clustered by experimental cohort rather than by genotype (**Fig.5C**). Consequently, we performed PCA on each experimental cohort independently to better resolve genotype-specific effects. The brain transcriptomes of MPS IIIA zebrafish appeared the most perturbed, forming distinct clusters from their wild-type siblings (**Fig. 5D,E**). Samples did not cluster by genotype in the MPS IIIB and MPS IIIC cohort (**Fig.5F**).

Differential gene expression analysis was subsequently employed to identify differentially expressed (DE) genes in each mutant relative to wild type controls adjusting for sex. Consistent with our PCA observations, the highest number of DE genes was detected in the MPS IIIA mutants (**Fig. 5G-I**, full output in **Supplemental Data File 2**). While MPS IIIA DE genes were the only set to show significant overrepresentation in GO terms, they were not consistent between experimental cohorts (**Fig.5J,K, S18**). Upon closer investigation of the DE genes in MPS IIIA zebrafish, we noticed that the DE genes in the MPS IIIA and MPS IIIB cohort were almost exclusively localised to chromosome 22, the same chromosome harbouring the *sgsh* mutation (with only 10 DEGs mapping elsewhere). While this chromosomal enrichment was observed across all mutations studied (**Fig.S19**), it was notably less severe in the MPS IIIA and MPS IIIC experimental cohort (where only 48 of 203 DE genes were on chromosome 22). This phenomenon likely reflects linkage disequilibrium rather than a systemic disease phenotype.

We next performed a *post-hoc* power calculation using *ssizeRNA* (Bi and Liu, 2016), which assesses the variation within the wild type group to quantify the statistical power at a given range of effect sizes. Consistent with the DE analyses, the lowest amount of statistical power was in the MPS IIIB and MPS IIIC cohort, where we achieved ∼60% power. Our two other cohorts reached ∼75% power to detect DE genes (**Fig.S20**). Given the reduced statistical power of the MPS IIIB and MPS IIIC cohort and the confounding chromosomal bias in the MPS IIIA and MPS IIIB cohort, we restricted our subsequent functional analyses to the MPS IIIA and MPS IIIC cohort.

In the MPS IIIA and MPS IIIC cohort, we detected significant over-representation of GO terms related to myelination and oligodendrocyte function in MPS IIIA zebrafish (**Fig.5K**,**S18**). No significant GO terms were enriched among the DE genes for MPS IIIC zebrafish. Recognising that over-representation analysis is limited by its reliance on a binary threshold for DE gene classification (e.g., FDR < 0.05), we employed the *fry* gene set test (Wu et al., 2010) to assess transcriptomic shifts more broadly. Unlike over-representation analysis, ***fry*** is a rotation-based, competitive gene set test that evaluates the collective behaviour of a gene set without requiring a hard significance cut-off for individual genes. To extend our analysis for MPS IIIB, given the insufficient statistical power of the current MPS IIIB cohorts, we re-analysed our previously published 6-month-old MPS IIIB RNA-seq dataset (Barthelson et al., 2025). While these fish represent a slightly later disease stage, we anticipated that the fundamental pathogenic pathways would remain consistent.

The KEGG gene sets significantly altered in at least one MPS III zebrafish brain are summarised in **Fig.6A**. There were fewer gene sets significantly altered in the MPS IIIC zebrafish than MPS IIIA and IIIB, consistent with MPS IIIC typically presenting with a milder clinical course (Lavery et al., 2017). Reassuringly, the KEGG gene sets altered in all MPS III zebrafish include *lysosome, glycosaminoglycan degradation*, and *other glycan degradation*. The significant changes to gene expression identified within these KEGG pathways are consistent with HS accumulation in lysosomes as the primary disease mechanism in MPS III, and with secondary accumulation of monosialic gangliosides (G_M2_ and G_M3_, reviewed in (Andrade et al., 2015)). These gene sets were broadly upregulated in all MPS III zebrafish brains (**Fig.6B-D**).

**Fig 6:**
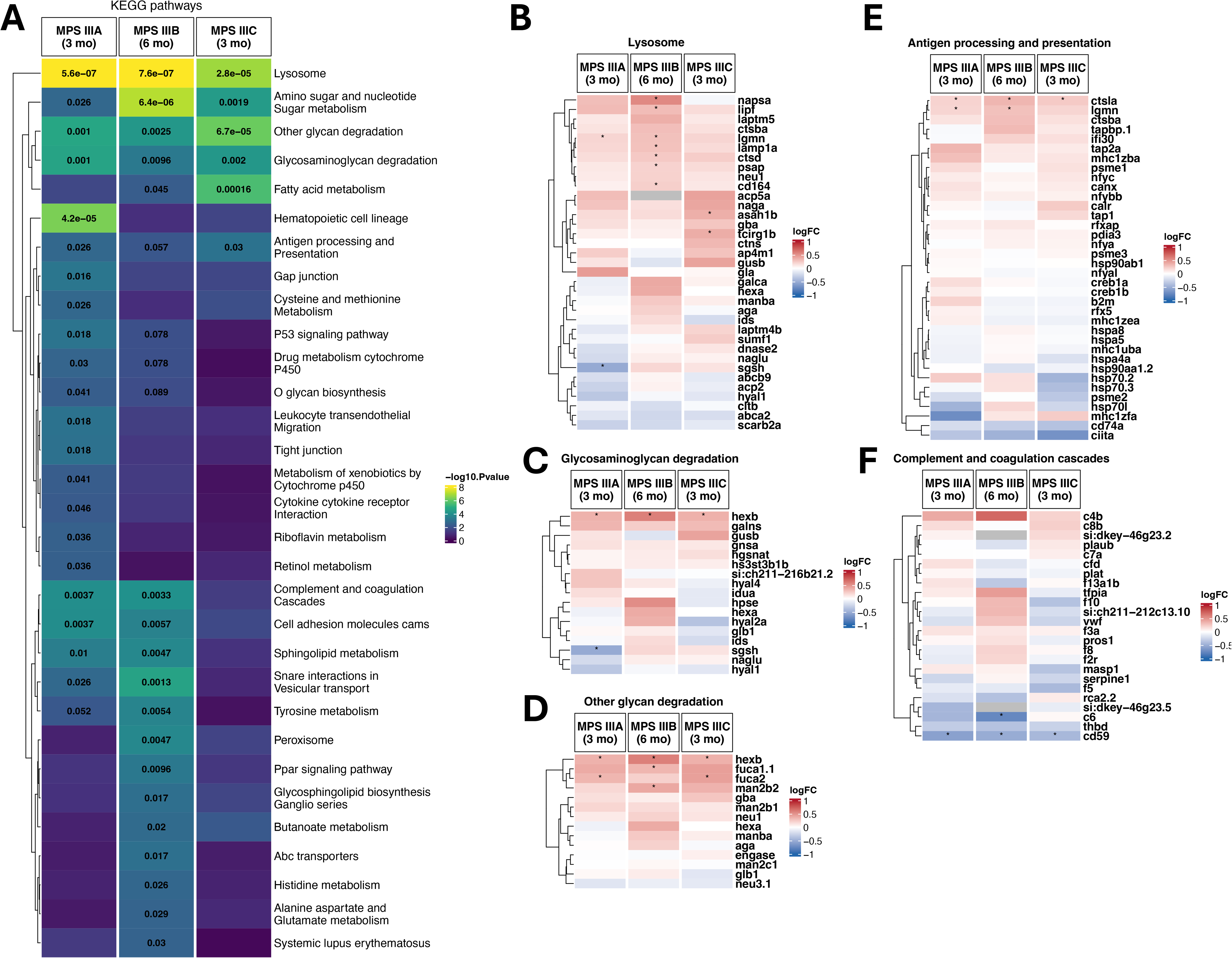
Differential expression of lysosomal, glycan degradation, immune, and complement pathway genes in adult MPS III mutant zebrafish brains. **A**, Heatmap indicating statistical significance in changes to gene expression in KEGG pathways. Only KEGG gene sets which reached the threshold of FDR-adjusted p-value of less than 0.05 in at least one MPS III zebrafish. The FDR-adjusted p-value (indicated within each cell) is displayed if it reached less than 0.1. **B-F**, Heatmap indicating the logFC values of genes in the **B**, lysosome, **C**, glycosaminoglycan degradation, **D**, other glycan degradation, **E** antigen processing and presentation, and **F**, complement and coagulation cascades gene sets in MPS III mutant zebrafish brains. Genes are labelled with an asterisk (*) if the FDR-adjusted p-value was less than 0.05 in differential expression testing.

Beyond primary metabolic defects, KEGG gene sets related to inflammatory processes were also altered. The *antigen presentation and processing* gene set was similarly dysregulated across all MPS III zebrafish transcriptomes, including upregulation of the cysteine proteases *ctsla, ctsba* and *lgmn* (**Fig.6E**). The *complement and coagulation cascades* gene set reached statistical significance only in the MPS IIIA and IIIB zebrafish brains. We observed a shared downregulation of *cd59*, a key regulator that inhibits membrane attack complex (MAC) assembly (reviewed in (Patel et al., 2024)) (**Fig.6F**). Its reduction suggests a vulnerability to MAC-mediated lytic pores, which induce osmotic and ion flux that can ultimately lead to cell death (reviewed in (Ho et al., 2025)). This dysregulation implies that CNS cells in our MPS III models may have altered vulnerability to terminal complement-mediated damage, reflecting a common neuroinflammatory baseline across the different genetic subtypes. Together, our functional analyses demonstrate that these zebrafish models recapitulate the core dysregulated pathways characteristic of MPS III pathophysiology.

### Changes to gene expression within neural stem cells, oligodendrocytes and microglia

The brain comprises multiple distinct cell types, and transcriptomic changes observed in bulk RNA-seq data may reflect shifts in cell-type proportions and/or altered gene expression within specific cell populations. To assess this, we defined cell-type marker gene sets using a published single-cell RNA-seq dataset of whole adult zebrafish brain (Tübingen strain, 4–12 months old, (Jiang et al., 2021)). Using *fry* with a directional hypothesis, we identified statistically significant overall reductions in expression of marker gene sets associated with oligodendrocytes, neural stem cells, and a macrophage population characterised by high *apoc1* expression (**Fig.7A**). The oligodendrocyte marker gene set showed widespread downregulation across all three mutant genotypes (**Fig.7B**). The macrophage marker gene set exhibited predominantly upregulated genes, with a weaker effect observed in MPS IIIC brains (**Fig.7C**). Marker genes associated with neural stem cells were also downregulated, although to a lesser extent than those of oligodendrocytes (**Fig.7D**). Notably, several DE genes in the neural stem cell gene set (*tspan2a, olig2, tuba8l3*, and *ninj2*) are also DE in the oligodendrocyte gene set. Whether these patterns reflect changes in cell-type abundance or altered transcriptional states within these populations cannot be resolved from bulk RNA-seq data alone and will require further investigation.

**Fig 7:**
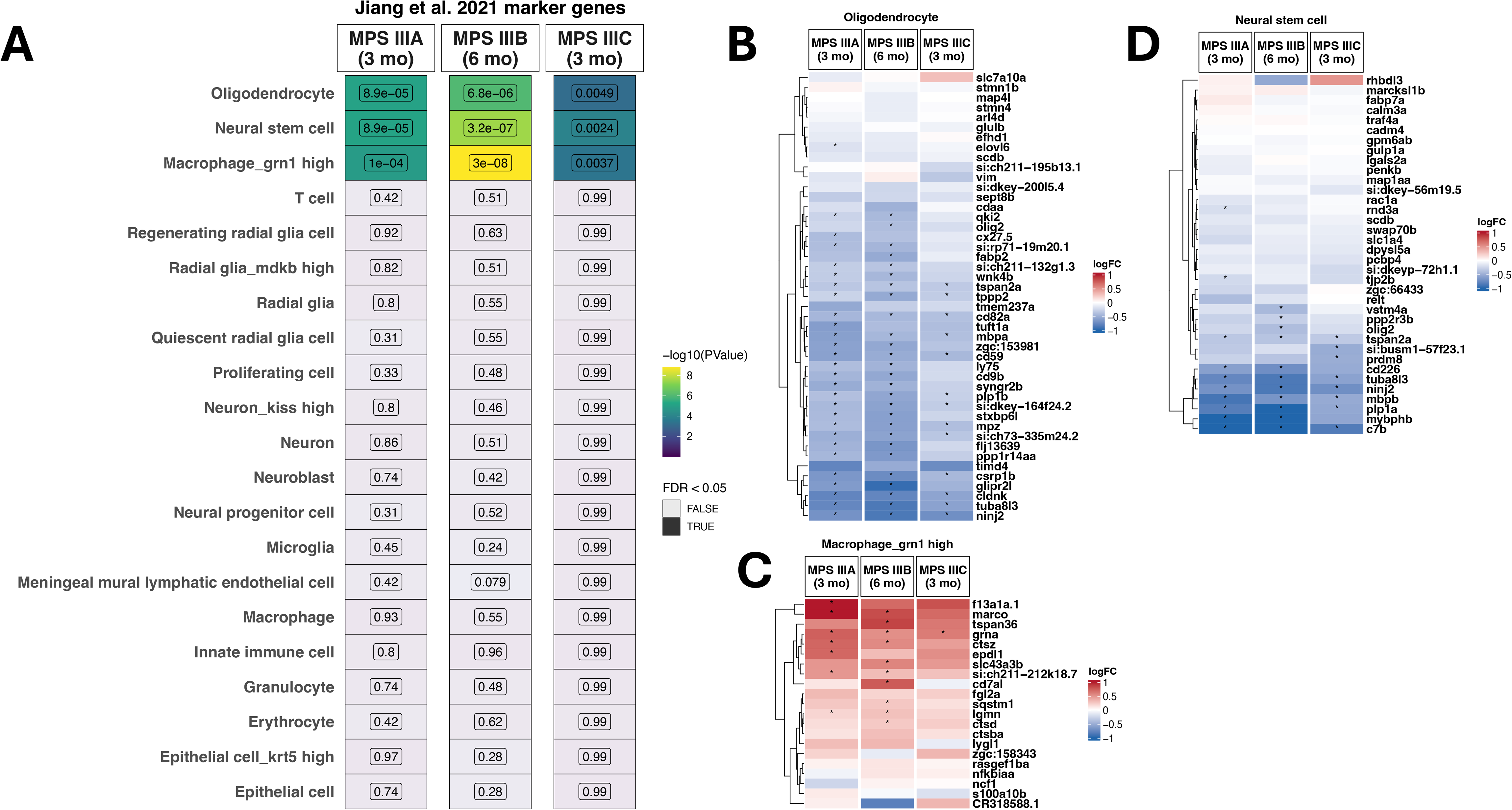
Directional changes to cell type marker gene expression in MPS III zebrafish brain transcriptomes. **A**, Heatmap indicating statistical significance in changes to gene expression in cell type marker gene sets from Jiang et al. 2021. Only KEGG gene sets which reached the threshold of FDR-adjusted p-value of less than 0.05 in at least one MPS III zebrafish. The FDR-adjusted p-value (indicated within each cell) is displayed if it reached less than 0.1. **B-D**, The log_2_FC (logFC) of detectable genes in the **B**, oligodendrocyte; **C**, macrophage; and **D**, neural stem cells gene sets. The colour represents the value of the logFC and significantly differentially expressed genes (FDR < 0.05) are marked with an asterisk (*).

### Evidence supporting iron dyshomeostasis in young MPS IIIA zebrafish brains

Changes to iron homeostasis are observed in many neurodegenerative diseases (reviewed in (Li and Reichmann, 2016)). Iron accumulation has been observed with deep brain MRI in human MPS IIIB cases (Brady et al., 2013) and with spectrometric analysis in the cortex mouse model of MPS IIIB (Puy et al., 2018). However, the broader role of iron homeostasis in all MPS III subtypes is poorly understood. To explore whether iron dyshomeostasis might occur in our zebrafish models of MPS III, we performed an enrichment analysis on our “IRE” gene sets (Hin et al., 2021). These gene sets consist of the genes in zebrafish encoding transcripts containing iron-responsive elements (IREs) in their 5’ or 3’ untranslated regions (UTRs). These gene sets also differentiate between those genes which encode consensus IRE sequences (those which have “high-quality” (ire_hq) IREs), and those with near-consensus sequences (the “ire_all” gene set) previously shown or predicted to function as IREs (Butt et al., 1996; Henderson et al., 1994). We identified significant changes to the expression of all IRE gene sets in only MPS IIIA zebrafish (**Fig.8**). The changes to the IRE genes were broadly similar across the other MPS III models, however, they did not reach statistical significance. Together, these findings suggest that changes to iron homeostasis are present in MPS IIIA zebrafish.

**Fig 8:**
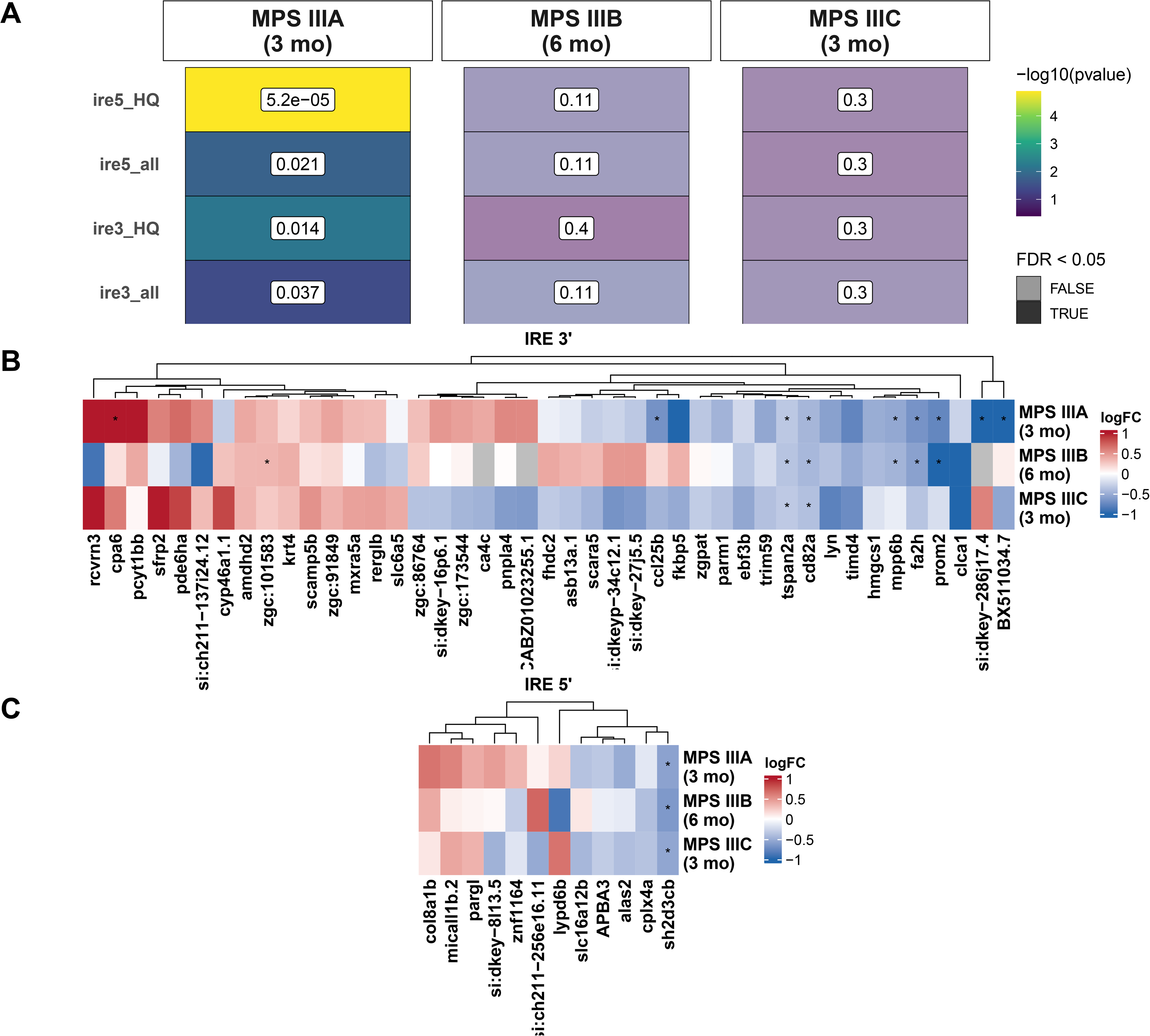
Iron homeostasis changes in MPS III zebrafish. **A**, Heatmap showing the statistical significance of the iron-responsive element (IRE) gene sets in MPS III zebrafish brains. The colour of the cells indicates the level of significance (brighter colour indicates greater statistical significance. The trans he transparency of the points (alpha) represents whether the FDR-adjusted p-value is > 0.05. Numbers within cells are the FDR-adjusted p-values. **B**, Heatmap indicating the log_2_FC (logFC) values for genes encoding transcripts with iron-responsive elements (IREs) in their 3’ or **C**, 5’ untranslated regions. Only genes showing absolute log_2_ fold change values greater than 0.3 are displayed for visualisation purposes, and significantly diffeentially expressed genes (FDR < 0.05) are labelled with an asterisk (*). Clustering was performed using Euclidean distances.

### No evidence that heterozygosity for a loss-of-function mutation in *sgsh* has effects on cellular functions in the brain

Single, heterozygous mutations in genes associated with other lysosomal storage disorders have been reported to be implicated in adult-onset neurodegenerative diseases (Clark et al., 2015; Gan-Or et al., 2013; Kluenemann et al., 2013). Recently, Douglass et al. (2021) identified subtle changes to motor coordination and brain pathology due to heterozygosity for a loss-of-function mutation in *Sgsh*. To investigate the molecular consequences of heterozygosity for a hypomorphic mutation in zebrafish *sgsh*, we analysed the brain transcriptomes of heterozygous *sgsh* ^*S387Lfs*^ fish in the MPS IIIA and MPS IIIB experimental cohort. A PCA performed on the brain transcriptome data from the MPS IIIA and MPS IIIB experimental cohort (and omitting the MPS IIIB zebrafish for clarity) revealed that the heterozygous *sgsh* ^*S387Lfs*^ fish appear to cluster with the wild type, rather than with MPS IIIA fish (**Fig.S21**), supporting that the heterozygous *sgsh* ^*S387Lfs*^ fish brain transcriptomes more closely represent a wild type state. Consistent with this observation, only 14 DE genes were detected in heterozygous *sgsh*^*S387Lfs*^ fish relative to wild type (**Fig.S21**), these were not significantly over-represented with any GO terms, and were mostly located on chromosome 22 (which contains *sgsh*). These chromosomally co-located differentially expressed genes (CC-DEGs) may reflect artefactual differential expression due to enrichment for linked expression quantitative trait loci (eQTLs), a phenomenon that can occur due to the breeding structure used to generate the families analysed (Baer et al., 2023; White et al., 2022). The DE genes in heterozygous *sgsh* ^*S387Lfs*^ fish mostly consisted of uncharacterised zebrafish transcripts without known function. Together, these observations do not support that zebrafish *sgsh*^*S387Lfs*^ mutation carriers show significant disturbances of brain function at 3 months of age.

## Discussion

Here, we described the creation and initial characterisation of novel zebrafish models of MPS IIIA, MPS IIIB and MPS IIIC. Together, our Sanger sequencing and *in silico* predictions suggest considerable changes to the primary, tertiary, and potentially quaternary, structures of each mutant protein. With this in mind, it should be noted that computationally predicted structures can bias toward folded states, so the mutations may affect folding and stability more severely *in vivo* (Buel and Walters, 2022). Consistent with these observations, we see highly significant accumulation of HS (**Fig.1D**), and reduction of enzyme activity (**Fig.3**) in the brains homozygous fish for all three mutants. The models thereby display the primary metabolic signature of the human disease. There was absence of any appreciable HS accumulation in zebrafish heterozygous for our mutations, consistent with the recessive nature of MPS III mutations in humans.

The *sgsh*^*S387Lfs*^ homozygous zebrafish generated and characterised here do not exhibit complete loss of sulfamidase activity, retaining approximately 3.6% of normal enzyme function. Despite this residual activity, substantial accumulation of HS was detected in the brains of homozygous mutants, indicating that this level of enzymatic function is insufficient to prevent substrate storage in the brain. Our MPS IIIA zebrafish model differs from the previously reported *sgsh*^*Δex5−6*^ model Douek et al. (2021), in which a large deletion spanning much of two exons and the joining intron results in complete loss of sulfamidase activity. Notably, our *sgsh*^*S387Lfs*^ mutation is similar to the human *SGSH* V379fs variant, which has been reported as one of the alleles in individuals with rapidly progressing MPS IIIA (Muschol et al., 2004; Trofimova et al., 2016; Wijburg et al., 2022). While enzyme activity associated with the human V379fs mutation has not been reported, a nearby frameshift mutation (*SGSH* V361Sfs), occurring within the same exon, shows no detectable sulfamidase activity in patient fibroblasts (Knottnerus et al., 2017). In addition, multiple nonsense and frameshift mutations associated with classical, early-onset MPS IIIA cluster in close proximity to this region of the protein (Yogalingam and Hopwood, 2001). Structural modelling of the mutant zebrafish Sgsh protein predicts disruption of regions critical for dimerisation, a process essential for sulfamidase function (Sidhu et al., 2014). We therefore hypothesise that the low level of residual enzymatic activity observed in homozygous mutants arises from inefficient or incomplete dimer formation. Nevertheless, despite partial preservation of enzymatic function, the pronounced substrate accumulation and disease-relevant phenotypes support the utility of this model for investigating the biological consequences of *SGSH* mutations.

Our *hgsnat*^*G577Sfs*^ mutation characterised here is analogous to the human *HGSNAT*^*T570Pfs*^ mutation. This human mutation has been reported as one of the alleles in an individual with *HGSNAT*-related non-syndromic retinitis pigmentosa, without clinical features of MPS IIIC (Schiff et al., 2020). In that patient, who carried a missense mutation on the second allele, residual HGSNAT enzyme activity was detected (0.3 nmol/hr/mg protein (Schiff et al., 2020)). In contrast, *hgsnat*^*G577Sfs*^ homozygous zebrafish exhibited no detectable enzyme activity, consistent with structural predictions indicating loss of two C-terminal transmembrane domains in the mutant protein. Our analysis suggests that these missing domains form critical dimerisation interfaces required for assembly of the active enzyme (Navratna et al., 2024). The discrepancy between residual enzyme activity reported in the human case and the complete loss observed in zebrafish is likely attributable to the presence of a second, missense allele in the patient, as enzyme activity associated with the *HGSNAT*^*T570Pfs*^ mutation in homozygosity has not been assessed. Intriguingly, despite the absence of detectable Hgsnat activity, *hgsnat*^*G577Sfs*^ homozygous fish exhibited the mildest transcriptional and behavioural alterations among the models examined. Together, these findings suggest that this mutation exerts relatively limited effects on the zebrafish brain at the ages studied and may therefore be more suitable as an attenuated model of MPS IIIC. Given the retinal phenotype associated with the equivalent human mutation, future studies should focus on characterising retinal structure and function in this model.

Our *naglu*^*A603Efs*^ mutation characterised here is most analogous to the human *NAGLU*^*E608Tfs*^ variant. While no clinical case reports have been published to date for this specific mutation, it is predicted to be pathogenic in ClinVar based on a single submission received (Accession: VCV001451924.5). Moreover, multiple frameshift mutations occurring downstream of the equivalent site in human *NAGLU* are associated with rapidly progressing MPS IIIB (Yogalingam and Hopwood, 2001), supporting the likelihood that disruption in this region of the protein is disease-relevant.

In contrast to the MPS IIIA and IIIC models, the *naglu*^*A603Efs*^ homozygous fish apparently exhibited 78% of wild type Naglu activity when assayed in 0.1% Triton X-100 buffer. Evaluation of Naglu activity where the *naglu*^*A603Efs*^ fish were instead homogenised in the enzyme buffer (0.2M sodium acetate pH 4.3) gave similar values (data not shown). Assuming structural homology with human NAGLU (Birrane et al., 2019), the *naglu*^*A603Efs*^ mutation is predicted to truncate domain III, which flanks the active-site cleft formed between domains II and III. One possibility is that the catalytic pocket of zebrafish Naglu (in contrast to human NAGLU) is not readily accessible to the artificial fluorogenic substrate (4-methylumbelliferyl-2-acetamido-2-deoxy-α-D-glucopyranoside) used in this assay. Alternatively, the mutation may impair correct lysosomal localisation of Naglu, such that enzymatic activity appears near-normal in a lysate-based assay despite functional deficiency *in vivo*.

Consistent with these interpretations, *naglu*^*A603Efs*^ homozygous fish exhibited the greatest fold increase in HS accumulation among the three models, exceeding that observed in age-matched *sgsh* ^S387Lfs^ and *hgsnat*^*G577Sfs*^ brains. This was accompanied by increased activities of wild-type Sgsh and Hgsnat enzymes, suggesting compensatory upregulation of adjacent lysosomal pathways. Together, these findings indicate a substantial defect in lysosomal Naglu function in our MPS IIIB zebrafish that is not adequately captured by the current biochemical assay.

In addition to biochemical abnormalities, *naglu*^*A603Efs*^ mutants displayed significant hyperactivity in the Y-maze and widespread changes in the brain transcriptome, including signatures beyond primary lysosomal dysfunction. Collectively, these molecular, biochemical, and behavioural phenotypes support the classification of the *naglu*^*A603Efs*^ zebrafish as a biologically relevant model of MPS IIIB.

Interestingly, all MPS III fish showed elevations in the activity of other lysosomal enzymes also involved in HS catabolism that were not directly related to the primary genetic defect. Similar observations have been made previously for a range of lysosomal hydrolases in an MPS IIIA mouse model (Bhaumik et al., 1999); our results are therefore not unexpected. These elevations are also congruent with the perturbations we observed in expression of lysosomal gene sets. This effect may be explained by an increase in lysosomal biogenesis via the upregulation of transcription factor EB (TFEB) caused by the lysosomal dysfunction exhibited in the affected animals (Sardiello et al., 2009). Of these off-target measured activities, Naglu activity in *hgsnat*^*G577Sfs*^ homozygous brains and Hgsnat activity in *sgsh*^*S387Lfs*^ homozygous brains were each not significantly elevated in our analysis. Since an obvious explanation for this (in the context of the HS degradative pathway) is lacking, it is possible that these activities are also elevated but to a more subtle extent not resolvable by this investigation’s analysis.

Zebrafish have great utility for behavioural studies due to their high rates of reproduction and short embryo development time. We employed intra-family behavioural analysis on our zebrafish models, since this is the best approach for reducing potentially confounding genetic and environmental effects. We also genotyped these fish after data collection. This has the benefits of preventing the results from being affected by genotyping procedure-imposed stress and of blinding the observer of the experiment. Our goal was to identify cognitive differences in our MPS III homozygous zebrafish. Cognitive functions, including memory deficits, can be difficult to assess accurately clinically in MPS III (Valstar et al., 2011). Children living with Sanfilippo syndrome are known to exhibit alterations in memory. However, to our knowledge, spatial working memory specifically has not been formally assessed in children with MPS III. However, spatial working memory differences have been observed in MPS IIIA mice (Crawley et al., 2006; Gliddon and Hopwood, 2004; Lau et al., 2025) and male MPS IIIB mice (Kan et al., 2016). We tested spatial working memory in our three zebrafish MPS III models but saw no significant differences between genotypes at the ages studied. Rather than reflecting any insensitivity of the Y-maze test to changes in spatial working memory in zebrafish compared to when these tests are conducted with mice and humans (Cleal et al., 2021b), the reason may lie in the remarkable neural regenerative abilities of zebrafish that contrast with those of mice and humans (reviewed by Zambusi and Ninkovic (2020)). Although we did not observe genotype-based differences in alternation, we did observe higher than normal proportions of repetition behaviour and lower than normal proportions of alternation across each MPS III zebrafish family. Previous work in this field has demonstrated that zebrafish, like humans and mice, employ an alternation-dominant search strategy in the Y-maze (Cleal et al., 2021b; Fontana et al., 2019). However, increased stress has been shown disrupt performance of zebrafish in similar spatial memory tests (Gaikwad et al., 2011). This suggests that our zebrafish may have been under additional stress due to unavoidable external stimuli, such as noise and vibration occurring during data collection, emphasising the need for follow-up experiments.

Hyperactivity is a well-established behavioural phenotype commonly displayed by children living with MPS III (Cleary and Wraith, 1993; Porter et al., 2021). We observed a significant hyperactivity phenotype in our aged MPS III *sgsh* and *naglu* model zebrafish but not in 5 dpf larvae. This is consistent with hyperactivity observed in mouse models of MPS IIIA (Hemsley and Hopwood, 2005; Langford-Smith, Alex et al., 2011) and MPS IIIB (Langford-Smith, A. et al., 2011) at widely varied ages. Although other mouse studies have produced conflicting results potentially related to environmental conditions and genetic background variation (Hemsley and Hopwood, 2005; Hemsley et al., 2007; Lau et al., 2008). While hyperactivity has been observed in MPS IIIC *Hgsnat* knock-out mice (Martins et al., 2015), we did not observe statistically significant hyperactivity in our *hgsnat* mutant zebrafish. This discrepancy may reflect species-specific differences in behavioural phenotypes, the generally milder clinical course of MPS IIIC in humans compared with other MPS III subtypes, and/or the nature of the zebrafish mutation studied here, which is analogous to a human mutation with a predominantly retinal phenotype. In humans with MPS III, hyperactivity is an early behavioural change preceding severe cognitive deficits (Cleary and Wraith, 1993) and so may be due to changes in, for example, synaptic activity before neurodegenerative cell loss (Dwyer et al., 2017; Vitry et al., 2009; Wilkinson et al., 2012). Therefore, in zebrafish models of MPS III with considerable regenerative ability, we might not expect to see changes in those behaviours due to cell loss, but might still observe them due to altered neuronal activity in largely intact brains. Therefore, additional testing of our MPS III models is warranted using behavioural tests for phenotypes not due to neurodegeneration. Differences in anxiety-related behaviours are common in human patients (Cleary and Wraith, 1993; Muschol et al., 2022), and have been observed in MPS IIIA (Lau et al., 2008), MPS IIIB (Cressant et al., 2004; Kan et al., 2016) and MPS IIIC (Martins et al., 2015) mice. While we assessed larval thigmotaxis in this study, other anxiety-related behaviours can also be assessed robustly in larval (Fontana and Parker, 2022) and adult (Fontana et al., 2022) zebrafish. Therefore these methods can be applied to our models in the future.

We aimed to implement a three-arm construction for our transcriptome analysis which, optimally, would enable direct comparison of each subtype to each other, as well as internal replication in comparing each subtype to wild type siblings twice. Unfortunately, the family structure and sample size achieved for the MPS IIIA and MPS IIIB, and the MPS IIIB and MPS IIIC cohorts meant we did not detect biologically meaningful changes to gene expression. Our analysis demonstrated the importance of appropriate sample size and of our intra-family approach, as zebrafish brain transcriptomes clustered very differently between parent pairs in the MPS IIIB and MPS IIIC cohort.

The MPS IIIA and MPS IIIC experimental cohort was our successful arm of the experiment, and we supplemented this dataset with our previously generated dataset containing MPS IIIB zebrafish at 6 months of age (Barthelson et al., 2025). Therefore, were thereby able to assess differential expression of KEGG pathways across the 3 MPS III subtypes. We detected differences in gene set enrichments in lysosomal and glycosaminoglycan degradation pathways, which are expected in the context of primary MPS III biology. We also detected increased expression of genes involved in inflammatory processes, consistent with increased complement protein expression and activated microglia phenotypes observed in other models of MPS III (Douek et al., 2021; Lau et al., 2025; Marcó et al., 2016; Ohmi et al., 2003).

The gene set analysis should be interpreted with a degree of caution given our observation of differential signals in the oligodendrocyte, macrophage and neural stem cell gene sets. We observed highly significant upregulation of a macrophage gene set. The differentially expressed genes in this gene set include a scavenger receptor (*marco*), lysosomal proteases (*ctsd, ctsz, lgmn*), and an autophagy-associated gene (*sqstm1*). This transcriptional profile is consistent with activated microglia (Sousa et al., 2018), and indicative of a neuroinflammatory response in the mutant brain.

The oligodendrocyte marker signal was consistently downregulated across all three models, suggesting either reduced transcriptional output within this lineage or a decreased abundance of mature oligodendrocytes. The changes to oligodendrocyte genes has been validated at the protein level in our MPS IIIB zebrafish at 6 months of age (Barthelson et al., 2025). One potential mechanism linking lysosomal pathology to this phenotype is excess HS, which is known to disrupt fibroblast growth factor (FGF) signalling by aberrant ligand sequestration and altered receptor engagement (De Pasquale et al., 2018; Wu et al., 2003; Yayon et al., 1991). FGF signalling plays a critical role in oligodendrocyte lineage progression by keeping oligodendrocyte precursor cells (OPCs) in a proliferative state, rather than maturation into oligodendrocytes (reviewed in (Bansal and Pfeiffer, 1997)). Consistent with this, reduced numbers of mature oligodendrocytes have been observed in symptomatic MPS IIIC mice (Taherzadeh et al., 2023). Loss of key oligodendrocyte proteins and ultrastructural abnormalities in myelin were also observed both in the mice, and in post-mortem MPS IIIC human brain tissue (Taherzadeh et al., 2023). The apparent sensitivity of oligodendrocytes in MPS III is biologically intuitive as the myelin sheathes of these cells give them the greatest surface area of any neural cell type and this is likely accompanied by the greatest burden of extracellular matrix (ECM) maintenance and turnover. This implies that oligodendrocytes would likely be more sensitive to deficits in degradation of HS (that is found abundantly in the ECM (reviewed in (Zhang et al., 2014)) than other cell types and would be the first to show pathological effects from mutations causing MPS III. Notably, oligodendrocytes are also the cell type showing the greatest accumulation of iron in the brain (Connor and Menzies, 1996; Reinert et al., 2019) and, if HS accumulation interferes with iron homeostasis (as our IRE analysis supported for MPS IIIA, discussed below), then oligodendrocytes would be expected to be particularly sensitive to this.

The gene set for neural stem cells (NSCs) was also observed to be downregulated. Although annotated as a NSC gene set, the differentially expressed genes within this gene set were enriched with genes associated with oligodendrocyte lineage specification and early maturation, including *olig2, plp1a, mbpb, tspan2a*, and *ninj2*. This suggests that the observed transcriptional changes within the NSC gene set may reflect disruption of oligodendrocyte differentiation trajectory rather than a global loss of NSCs.

We observed significant alterations in the expression of genes encoding transcripts that contain iron-responsive elements (IREs) within their untranslated regions specifically in MPS IIIA zebrafish (and not MPS IIIB or IIIC). Traditionally, it was thought binding of an iron-responsive element-binding protein (IRP) to an IRE in a transcript’s 3’UTR would stabilise that transcript, while IRP binding to a 5’ IRE would block translation of the transcript. However, Hin et al.’s study of changes in IRE-containing transcript levels under iron deficient and iron excess conditions showed that the particular direction of level change of any transcript was not a general consequence of any IRE type (5’ or 3’ UTR). Therefore, the IRE changes observed cannot be used to impute iron deficiency or excess in MPS IIIA brains. The acidic internal environment of the lysosome is required for the reduction of the biologically inactive ferric form of iron (Fe^3+^) to the active ferrous form of iron (Fe^2+^) (Lambe et al., 2009; Yambire et al., 2019). When HS accumulates in lysosomes, it may upset the normal acidification of these compartments and interfere with the proper conversion of ferric to ferrous iron. This would have significant implications for neural cells, given the central role of ferrous iron in energy metabolism, the dependence of oligodendrocytes on iron and other crucial molecular processes.

MPS III is devastating for children living with the disease and their families. With limited therapeutic options available, animal models are a crucial part of efforts to understand and treat the underlying pathologies. In this study, we successfully generated novel zebrafish models of MPS IIIA, MPS IIIB and MPS IIIC by creating the hypomorphic mutations *sgsh*^*S387Lfs*^, *naglu*^*A603Efs*^ and *hgsnat*^*G577Sfs*^. To our knowledge, our MPS IIIB *naglu* and MPS IIIC *hgsnat* models are the first zebrafish models of these disorders. Importantly, these models feature significant HS accumulation in the brain, displaying the primary signature of the disease. We found that, in contrast to the already published zebrafish mutation, *sgsh*^*Δex5−6*^, our *sgsh*^*S387Lfs*^ mutation appears to retain some enzyme function, a feature which could be useful in future investigations of this heterogeneous disease. While zebrafish intra-family behavioural analysis did not provide evidence of effects on adult memory or larval behaviour, two of our models displayed statistically significant hyperactivity in the homozygous state at later ages, a phenotype in-common with MPS III in humans. Intra-family RNA-seq transcriptome analysis revealed expected changes in gene expression related to lysosomal and glycan-degradative biology and the immune system in MPS III zebrafish brains. Interestingly, we also observed evidence suggesting iron dyshomeostasis in MPS IIIA zebrafish, and changes in oligodendrocyte cell state and neuroinflammation in all three mutants. Ultimately, our investigations of the behavioural and molecular features of our MPS III zebrafish show concordance both with existing mouse models and the human condition. Further investigation is required to deepen our understanding of the molecular pathways involved and for replication of our findings. We envisage that the *sgsh*^*S387Lfs*^, *naglu*^*A603Efs*^ and *hgsnat*^*G577Sfs*^ mutant zebrafish will be valuable platforms for discovery and testing of novel MPS IIII therapeutic approaches.

## Acknowledgements

The authors of this paper would like to express their sincere appreciation to several individuals who made valuable contributions to this project. We thank Dr. Morgan Newman and Rebel Bertram for their assistance with the initial screening for generation of the mutants, Barb King for her invaluable assistance with fluorometric enzyme assays, Associate Professor Marten Snel and Dr Paul Trim (SAHMRI) for performing the HS measurements, Dr Matt Parker for advice regarding behaviour analysis, the South Australian Genomics Centre (SAGC) for performing the library preparation and RNA-sequencing, Shijin Suo for preliminary experiments and Dr. Lachlan Baer for help with the *snakemake* pipeline.

## Author Contributions

ML was responsible for project conception and funding. ML and KB were responsible for oversight. EG, KB and SA generated and characterised the mutants. AA performed the *in silico* protein modelling. For the enzyme activity assays, EG collected the samples and AL performed the assays. EG and LR generated and analysed the adult and larval behaviour data respectively. LR was supervised by KH and KB. KB and EG generated the RNA-seq data, which was then analysed by KB. Initial drafting of the manuscript was performed by EG and KB. Subsequent revisions and editing were performed by all authors.

