## Supplemental Figures and Tables for "Zebrafish models of Mucopolysaccharidosis IIIA, IIIB, and IIIC show hyperactivity, neuroinflammation and changes in oligodendrocyte cell state"

**Table S1. CRISPR sequences used for mutagenesis of zebrafish orthologues of MPS III-involved genes**

| **Gene** | **Location** | **Strand** | **crRNA sequence** | **PAM** |
| --- | --- | --- | --- | --- |
| *sgsh* | Exon 7 | + | CCATCCACAAGGGCCCCTAC | CGG |
| *naglu* | Exon 7 | + | TGACCTCCTCACGGCTGGCG | TTTC |
| *hgsnat* | Exon 17 | + | GATGTAAAGAAGTGGTGGTC | TGG |

**Table S2. Primers used for the PCR amplification of genomic DNA for T7 endonuclease assays, Sanger sequencing, and family genotyping**

| **MPS III Genes** | **Primers** | **Sequence (5’ – 3’)** | **T_m_ (ºC)** | **T_a_ (ºC)** | **Extension time (s)** | **Amplicon length (bp)** | **Purpose** |
| --- | --- | --- | --- | --- | --- | --- | --- |
| *sgsh* | Forward | GTCGTCCCTCACTATTGTTTCCG | 67.5 | 60 | 60 | 702 | T7 assay |
|  | Reverse | CGTCATGTTTTTGTTTTTATTCAGA | 62.7 |  |  |  |  |
| *naglu* | Forward | AAGATAGAATGTCAGTCAAAGTGCT | 61.1 | 60 | 60 | 1199 | T7 assay |
|  | Reverse | TCCTGTGGTTTGTAATGTAATGGG | 65.3 |  |  |  |  |
| *hgsnat* | Forward | GCATTTGTCTCTTGTTTTAGGTCT | 61.4 | 60 | 60 | 641 | T7 assay |
|  | Reverse | CAAACTCCACACAGAAACGCTAAC | 65.6 |  |  |  |  |
| *sgsh* | Forward | GTCGTCCCTCACTATTGTTTCCG | 67.5 | 65 | 30 | 334 | Sanger seq & genotyping |
|  | Reverse | TCGTCCTGACTGAGTGCGGT | 68.6 |  |  |  |  |
| *naglu* | Forward | AGAGATATTCACAGACTGTTGCC | 61.1 | 60 | 30 | 224 | Sanger seq & genotyping |
|  | Reverse | TTGCTGGAAAGGATGCGG | 67.3 |  |  |  |  |
| *hgsnat* | Forward | GCCATCAGAAGATTCCAACACA | 66.1 | 60 | 60 | 745 | Sanger seq & genotyping |
|  | Reverse | TGCCCTAGTACACATTAAGCCT | 61.5 |  |  |  |  |

**Table S3. Welch ANOVA test values for effect of *sgsh^S387Lfs^*, *naglu^A603Efs^* and *hgsnat^G577Sfs^* genotype on enzyme activities and substrate accumulation.** DFn: Degrees of freedom, numerator, DFd: Degrees of freedom, denominator

| Sgsh activity | | | | | |
| --- | --- | --- | --- | --- | --- |
| **Mutation** | **n** | **F statistic** | **DFn** | **DFd** | **p-value** |
| *sgsh^S387Lfs^* | 15 | 88.74 | 2 | 5.89 | 3.98E-05 |
| *naglu^A603Efs^* | 15 | 30.63 | 2 | 6.94 | 0.000359 |
| *hgsnat^G577Sfs^* | 15 | 42.18 | 2 | 7.62 | 7.54E-05 |
| Naglu activity | | | | | |
| **Mutation** | **n** | **F statistic** | **DFn** | **DFd** | **p-value** |
| *sgsh^S387Lfs^* | 15 | 62.05 | 2 | 7.67 | 1.83E-05 |
| *naglu^A603Efs^* | 15 | 10.91 | 2 | 7.27 | 0.006 |
| *hgsnat^G577Sfs^* | 15 | 3.21 | 2 | 6.7 | 0.105 |
| Hgsnat activity | | | | | |
| **Mutation** | **n** | **F statistic** | **DFn** | **DFd** | **p-value** |
| *sgsh^S387Lfs^* | 15 | 2.03 | 2 | 7.65 | 0.196 |
| *naglu^A603Efs^* | 15 | 36.47 | 2 | 7.59 | 0.000128 |
| *hgsnat^G577Sfs^* | 15 | 74.27 | 2 | 5.35 | 0.000125 |
| HS accumulation | | | | | |
| **Mutation** | **n** | **F statistic** | **DFn** | **DFd** | **p-value** |
| *sgsh^S387Lfs^* | 9 | 15.77 | 2 | 3.176196 | 0.022 |
| *naglu^A603Efs^* | 9 | 167.7 | 2 | 2.697887 | 0.001 |
| *hgsnat^G577Sfs^* | 9 | 14.67 | 2 | 3.434771 | 0.021 |

**Table S4: Type II Wald 𝝌2 test values for generalised linear model analysis of total turns performed by MPS III adult zebrafish families in the Y-maze.** Df:degrees of freedom

| ***sgsh^S387Lfs^*** | | | | ***naglu^A603Efs^*** | | | | ***hgsnat^G577Sfs^*** | | | |
| --- | --- | --- | --- | --- | --- | --- | --- | --- | --- | --- | --- |
|  | *𝝌2* | *Df* | *p-value* |  | *𝝌2* | *Df* | *p-value)* |  | *𝝌2* | *Df* | *p-value* |
| **genotype** | 7.54 | 2 | 0.023 | **genotype** | 13.8 | 2 | 0.001 | **genotype** | 1.22 | 2 | 0.543 |
| **bin** | 243 | 5 | 1.83E-50 | **bin** | 227 | 5 | 4.60E-47 | **bin** | 128 | 5 | 5.70E-26 |
| **tank** | 1.92 | 3 | 0.589 | **tank** | 1.16 | 3 | 0.762 | **tank** | 2.28 | 3 | 0.515 |
| **sex** | 0.142 | 1 | 0.706 | **sex** | 0.211 | 1 | 0.646 | **sex** | 0.876 | 1 | 0.349 |
| **geno:bin** | 26.6 | 10 | 0.003 | **geno:bin** | 1.91 | 10 | 0.997 | **geno:bin** | 9.71 | 10 | 0.466 |

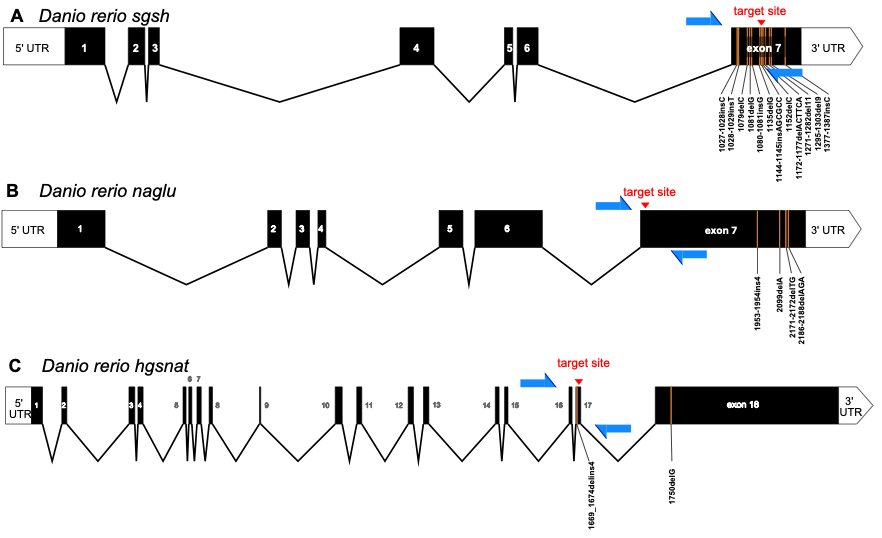

**Fig.S1: Exon-intron structure of *Danio rerio* A) *sgsh*, B) *naglu* and C) *hgsnat* showing mutagenesis targeting strategy.** Target sites (red) were chosen based on their proximity to known MPS III-causative mutations in the equivalent human genes (orange) (representative only, not to scale, MPS III-causative mutations in other exons are not shown). Positions of genomic DNA amplification PCR primer binding are indicated in blue (not to scale). Human mutations were soured from Yogalingam and Hopwood (2001) for *SGSH* and *NAGLU* mutations, and Feldhammer et al. (2009) for *HGSNAT*.

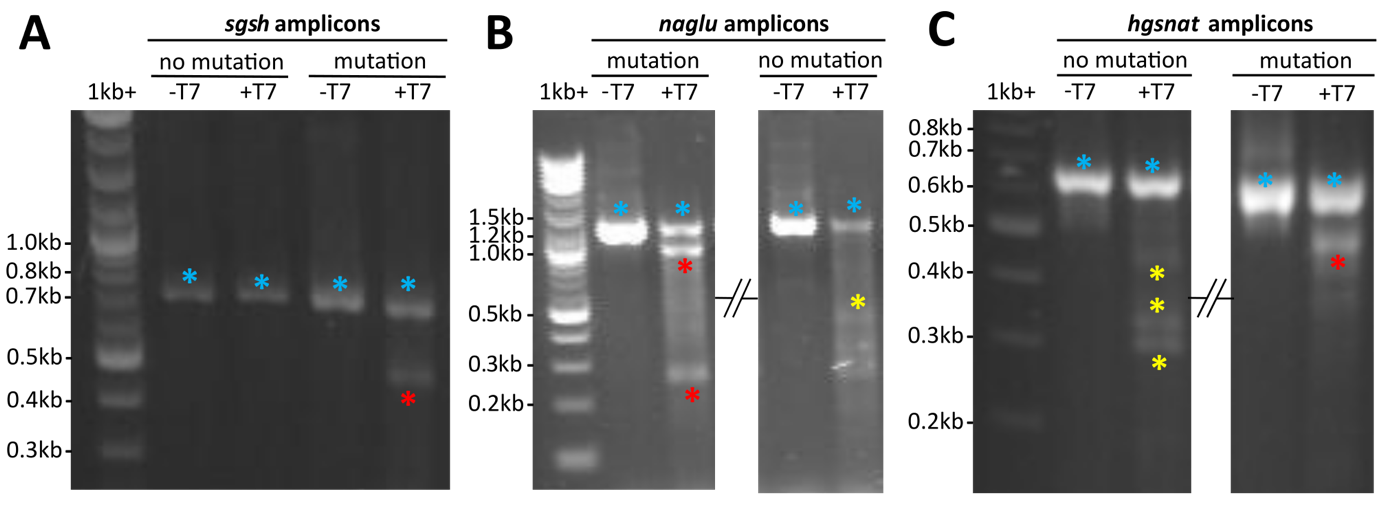

**Fig.S2: Mutation screening by T7 endonuclease reactions in Tübingen zebrafish following targeted CRISPR mutagenesis.** Shown are images from agarose gel electrophoresis of PCR amplicons produced using primers (**Table S2**) surrounding the mutation target sites of **A,** *sgsh*; **B,** *naglu***;** and **C,** *hgsnat*. For each gene, a sample with an on-target mutation and a sample with no mutation of interest are shown. Each sample is divided into a negative control aliquot electrophoresed adjacent to an aliquot after incubation with T7 endonuclease. Bands are specified with coloured asterisks; undigested amplicons (blue), cleavage fragments at sizes expected given the presence of a targeted mutation (red), and non-specific cleavage fragments (yellow). *sgsh*: 702 bp cleaved to ~480 bp and 220 bp, *naglu*: 1199 bp cleaved to ~950 bp and 250 bp and *hgsnat*: 641 bp cleaved to ~520 bp and 120 bp. Approximate DNA fragment sizes are indicated corresponding to the 1kb+ ladder (NEB®). In **B** and **C**, a section of the gel has been removed for visual concision.

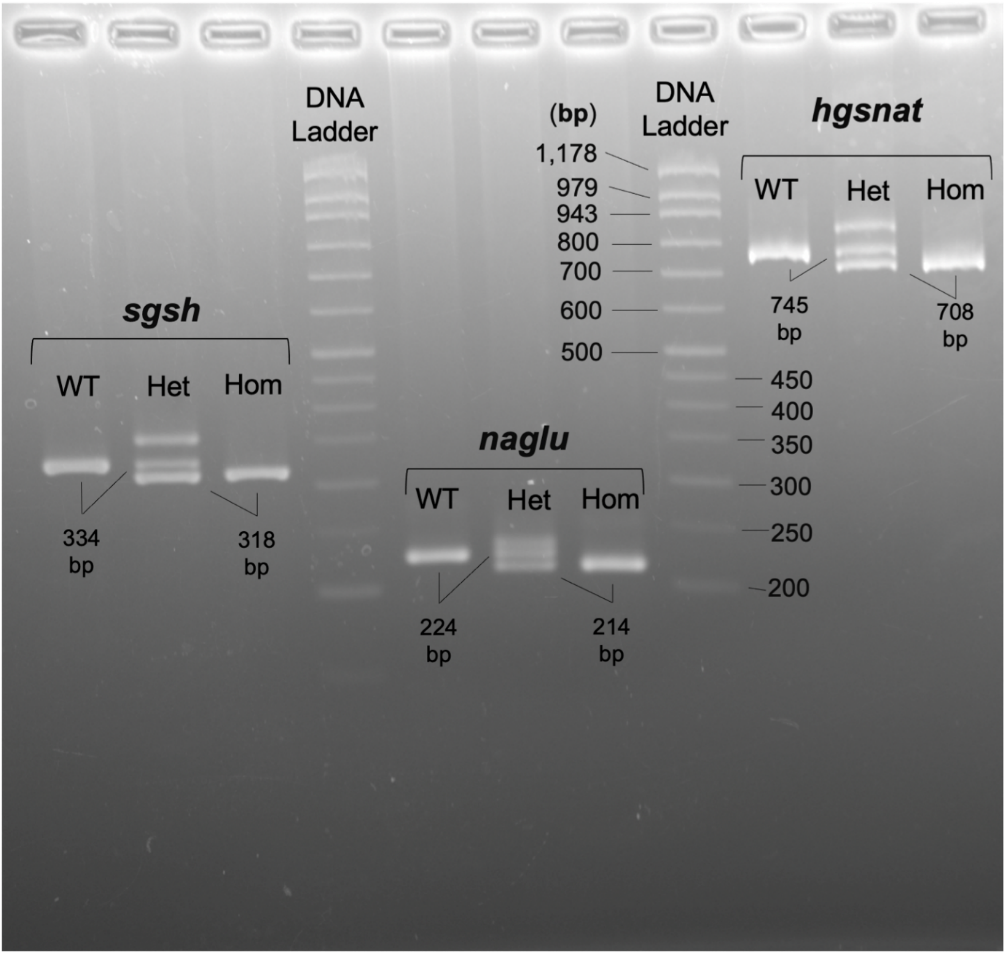

**Fig.S3: Example agarose gel electrophoretograms for genotyping of MPS III model zebrafish.** From left to right, lanes 1-3 contain PCR products containing amplified fragments of *sgsh* DNA from larvae that are wild type (WT), heterozygous (Het) and homozygous (Hom) for the *sgsh^S387Lfs^* mutation. Expected product size for the WTs is 334 bp, the Homs are expected to be 308 bp and Hets are expected to have both DNA fragments. Lanes 5-7 display amplified DNA bands containing a fragment of the *naglu* gene from larvae WT (224 bp), Het (224 bp and 214 bp) and Hom (214 bp) for the *naglu^A603Efs^* mutation. Lanes 9-11 are comprised of amplified fragments of the *hgsnat* gene from samples that are WT (745 bp), Het (745 bp and 708 bp) and Hom (708 bp) for the *hgsnat^G577Sfs^* mutation. pHAPE BamHI DNA ladder [24] was used to determine the size of the bands. The third band for all heterozygous samples is due to the formation of heteroduplexes between wild type and mutant fragments.

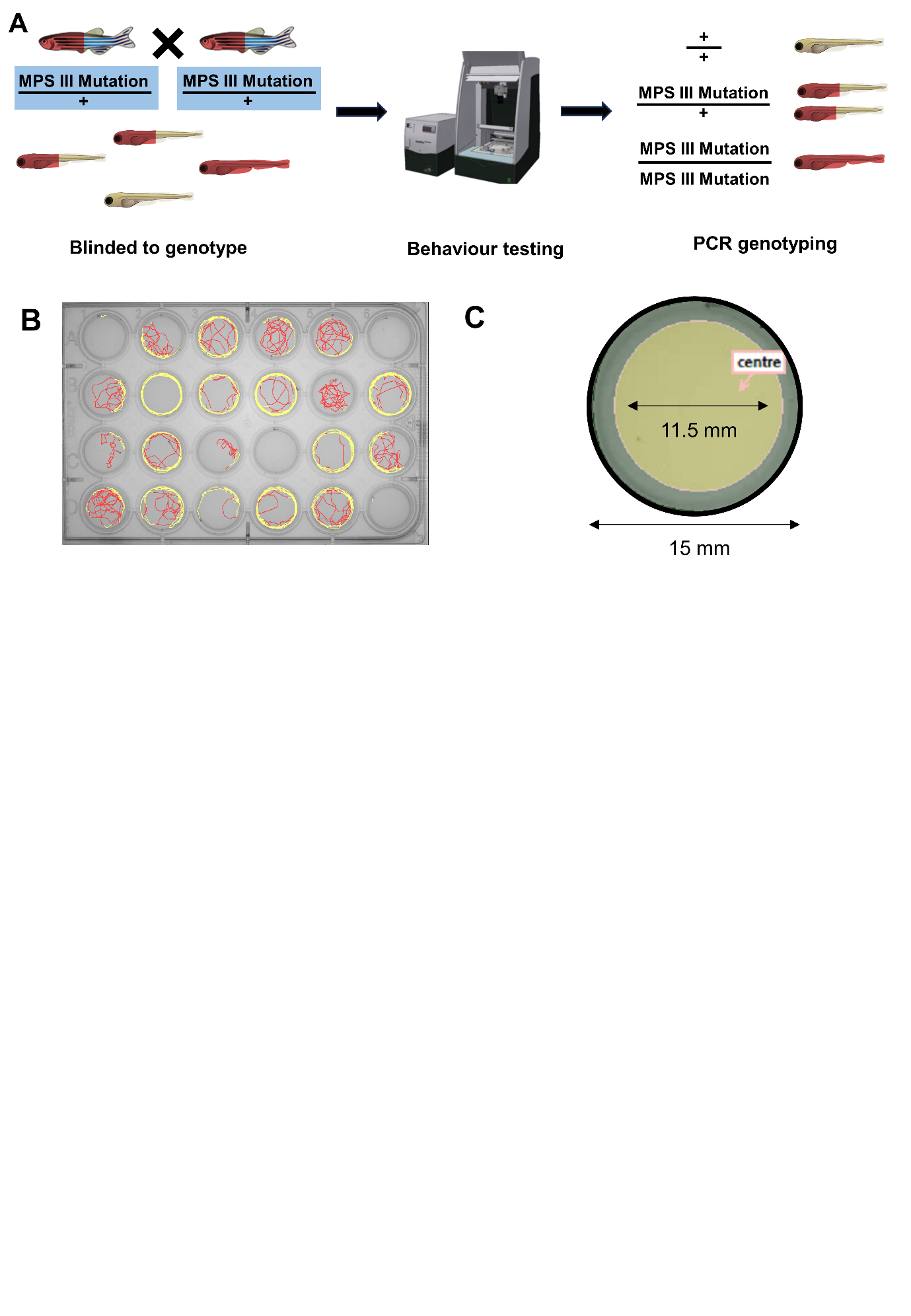

**Fig.S4: Experimental design for behavioural testing of zebrafish larvae. A,** Pairs of zebrafish heterozygous for either *sgsh^S387Lfs^, naglu^A603Efs^* or *hgsnat^G557Sfs^* are in-crossed, resulting in families of progeny with the expected ratio of 1:2:1 for wild type, heterozygous and homozygous genotypes respectively. **B**, Larvae are raised together before separation of individuals into the individual wells of 24-well plates and tracked in a DanioVision Observation Chamber (Noldus) for one hour. Panel **C** depicts the dimensions of a single well with central and outer zones. Larvae are genotyped following behavioural testing using PCR and gel electrophoresis.

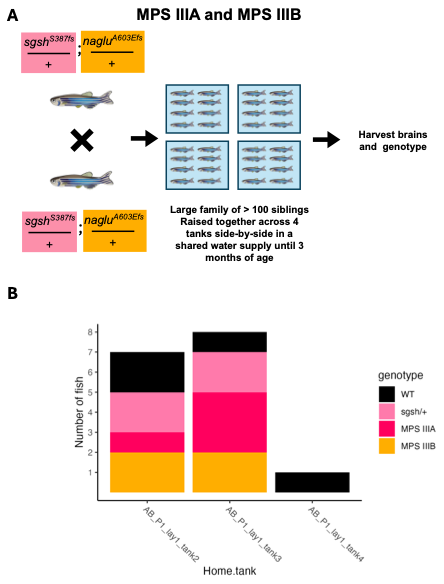

**Fig.S5: Detailed study design for the MPS IIIA and MPS IIIB experimental cohort. A,** Zebrafish doubly heterozygous for the MPS IIIA and MPS IIIB mutations (*sgsh^S387Lfs^* and *naglu^A603Efs^* respectively) are in-crossed, to generate a large family of at least 100 siblings. This large family is raised across 4 tanks, at a density of 25 fish per tank until 3 months of age. Then, all fish in the family are euthanised and their brains removed and preserved in RNA*later* solution. Each fish is genotyped at the *sgsh^S387Lfs^* and *naglu^A603Efs^* sites by PCRs. **B,** Number of fish per tank analysed in the RNA-seq experiment.

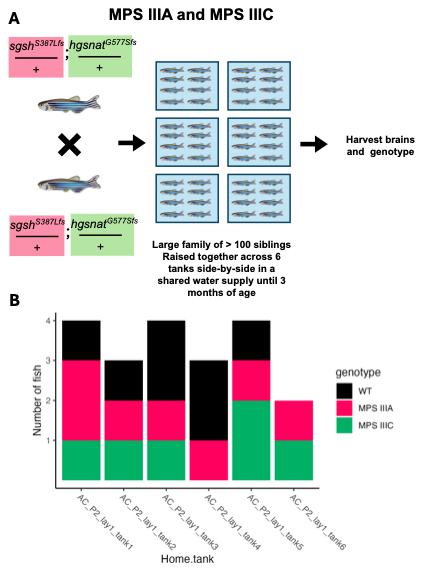

**Fig.S6: Detailed study design for the MPS IIIA and MPS IIIC experimental cohort. A,** Zebrafish doubly heterozygous for the MPS IIIA and MPS IIIC mutations (*sgsh^S387LFS^* and *hgsnat^G577Sfs^* respectively) are in-crossed, to generate a large family of at least 100 siblings. This large family is raised across at least 4 tanks, at a density of 25 fish per tank until 3 months of age. Then, all fish in the family are euthanised and their brains removed and preserved in RNA*later* solution. Each fish is genotyped at the *sgsh^S387Lfs^*  and *hgsnat^G577Sfs^* sites by PCRs. **B,** Number of fish per tank analysed in the RNA-seq experiment.

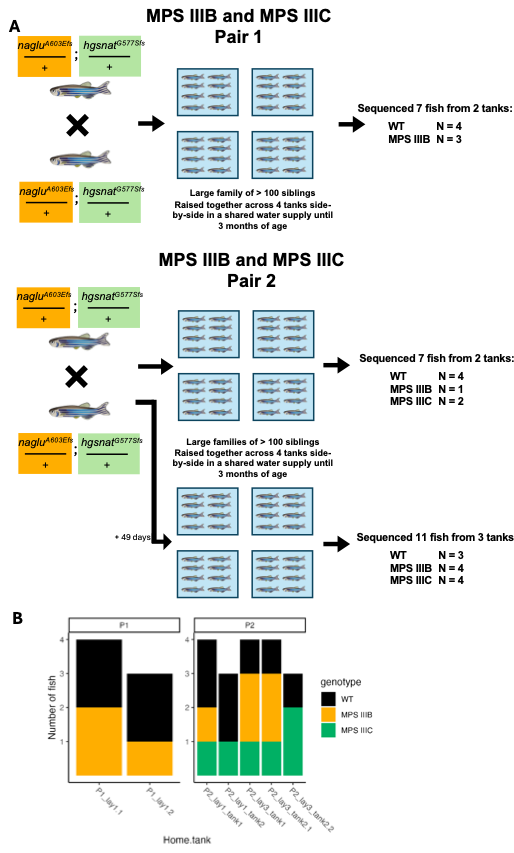

**Fig.S7: Detailed study design for the MPS IIIB and MPS IIIC experimental cohort. A,** Two pairs (P1 and P2) of zebrafish doubly heterozygous for the MPS IIIB and MPS IIIC mutations (*naglu^A603Efs^* and *hgsnat^G577Sfs^* respectively) were in-crossed, to generate large families of at least 100 siblings. These large families was raised across 8 tanks, at a density of 25 fish per tank until 3 months of age. Then, all fish in the families were euthanised and their brains removed and preserved in RNA*later* solution. Each fish is genotyped at the *naglu^A603Efs^* and *hgsnat^G577Sfs^* sites by PCRs. Insufficient fish per genotype were identified. Therefore, we included another family of fish at 3 months of age which arose from the P2 parents and which were spawned 49 days after the first clutch. **B,** Number of fish per tank analysed in the RNA-seq experiment.

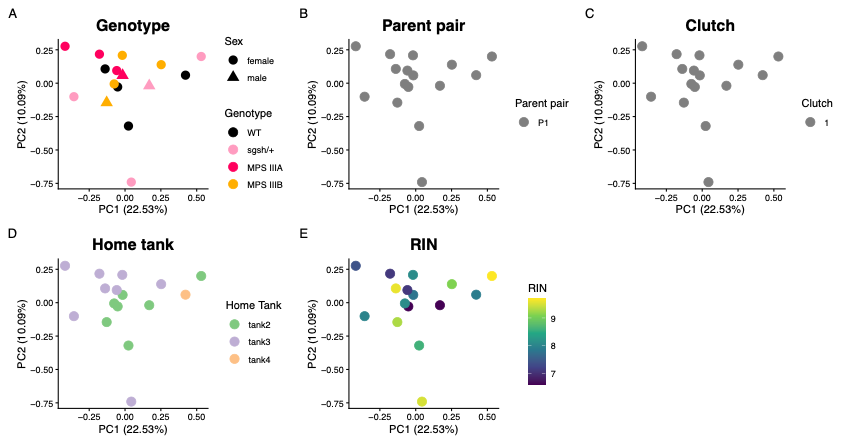

**Fig.S8: Principal component analysis on the MPS IIIA and MPS IIIB cohort.** Each point represents a brain transcriptome which is coloured according to **A,** MPS III genotype; **B,** the parental pair from which the progeny fish was spawned; **C,** the particular clutch to which the fish belonged; **D,** the tank in which the fish was raised; and **E,** the RNA integrity number (RIN) indicating the quality of the brain RNA sequenced.

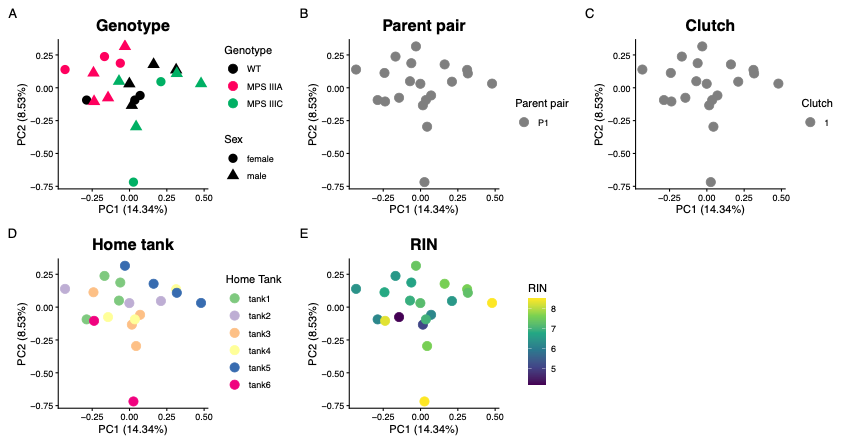

**Fig.S9: Principal component analysis on the MPS IIIA and MPS IIIC cohort.** Each point represents a brain transcriptome which is coloured according to **A,** MPS III genotype; **B,** the parental pair from which the progeny fish was spawned; **C,** the particular clutch to which the fish belonged; **D,** the tank in which the fish was raised; and **E,** the RNA integrity number (RIN) indicating the quality of the brain RNA sequenced.

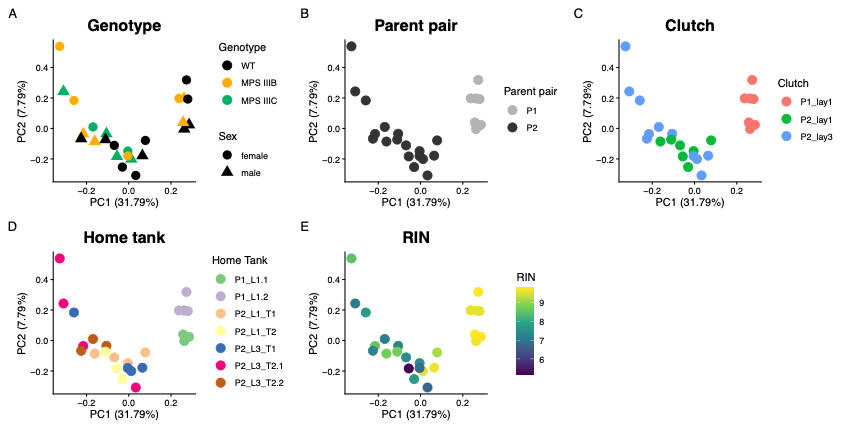

**Fig.S10: Principal component analysis on the MPS IIIB and MPS IIIC cohort.** Each point represents a brain transcriptome which is coloured by **A,** MPS III genotype; **B,** the parental pair from which the progeny fish was spawned; **C,** the particular clutch to which the fish belonged; **D**, the tank in which the fish was raised; and **E,** the RNA integrity number (RIN) indicating the quality of the brain RNA sequenced.

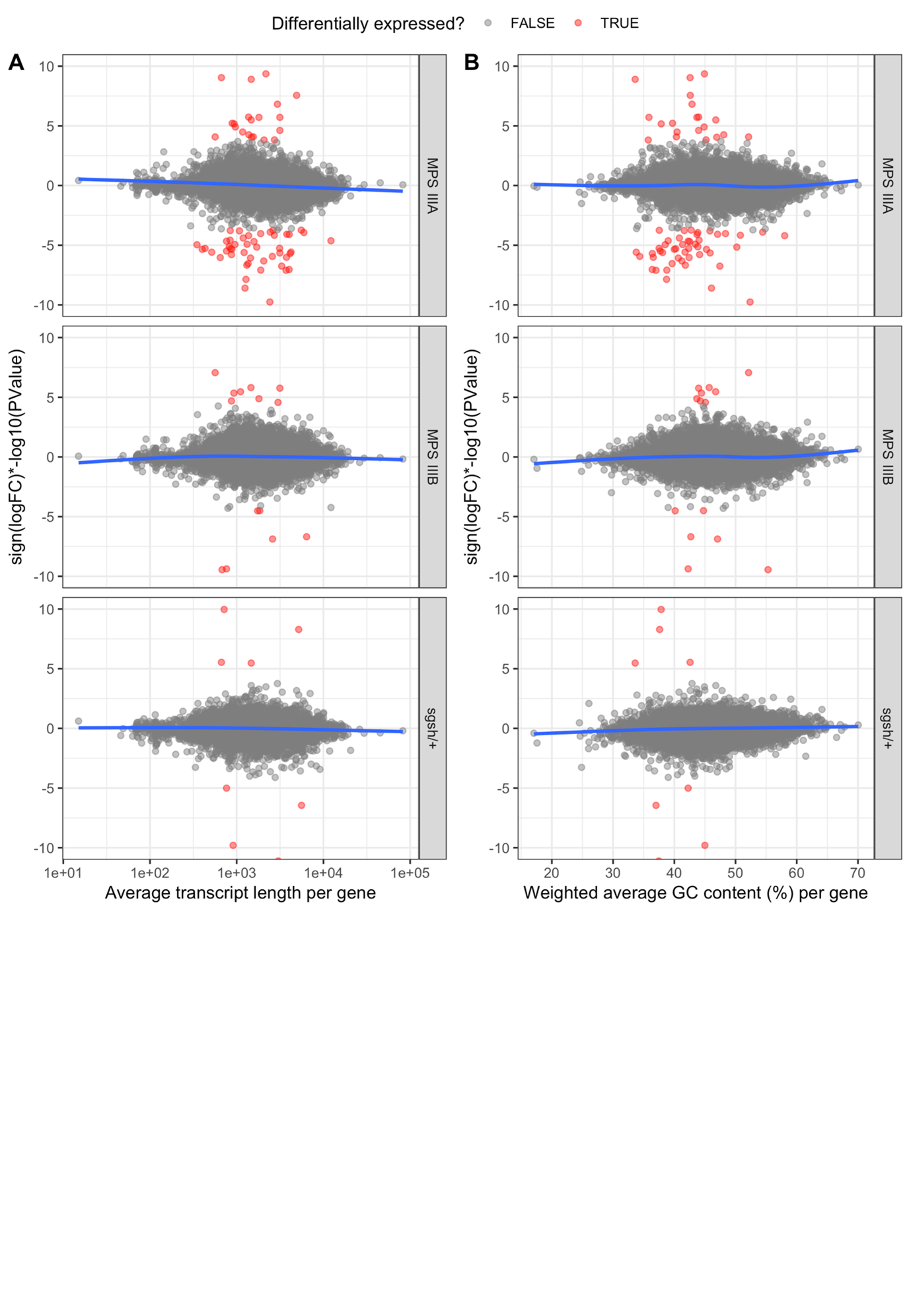

**Fig.S11:** **Observed bias for differential expression with GC content and length in the MPS IIIA and MPS IIIB cohort**. A ranking metric was calculated using the sign of logFC multiplied by − log_10_ of the p-value against **A,** the weighted (by transcript length) %GC content per gene; and **B,** the average transcript length per gene. The blue curve indicates the line of best fit from a generalised additive model. Given the gam fit is a not completely horizontal line mostly overlapping y = 0, a small bias for %GC content or length is likely present in this dataset. The ranking statistic limits were constrained to − 10 and 10 for visualisation purposes.

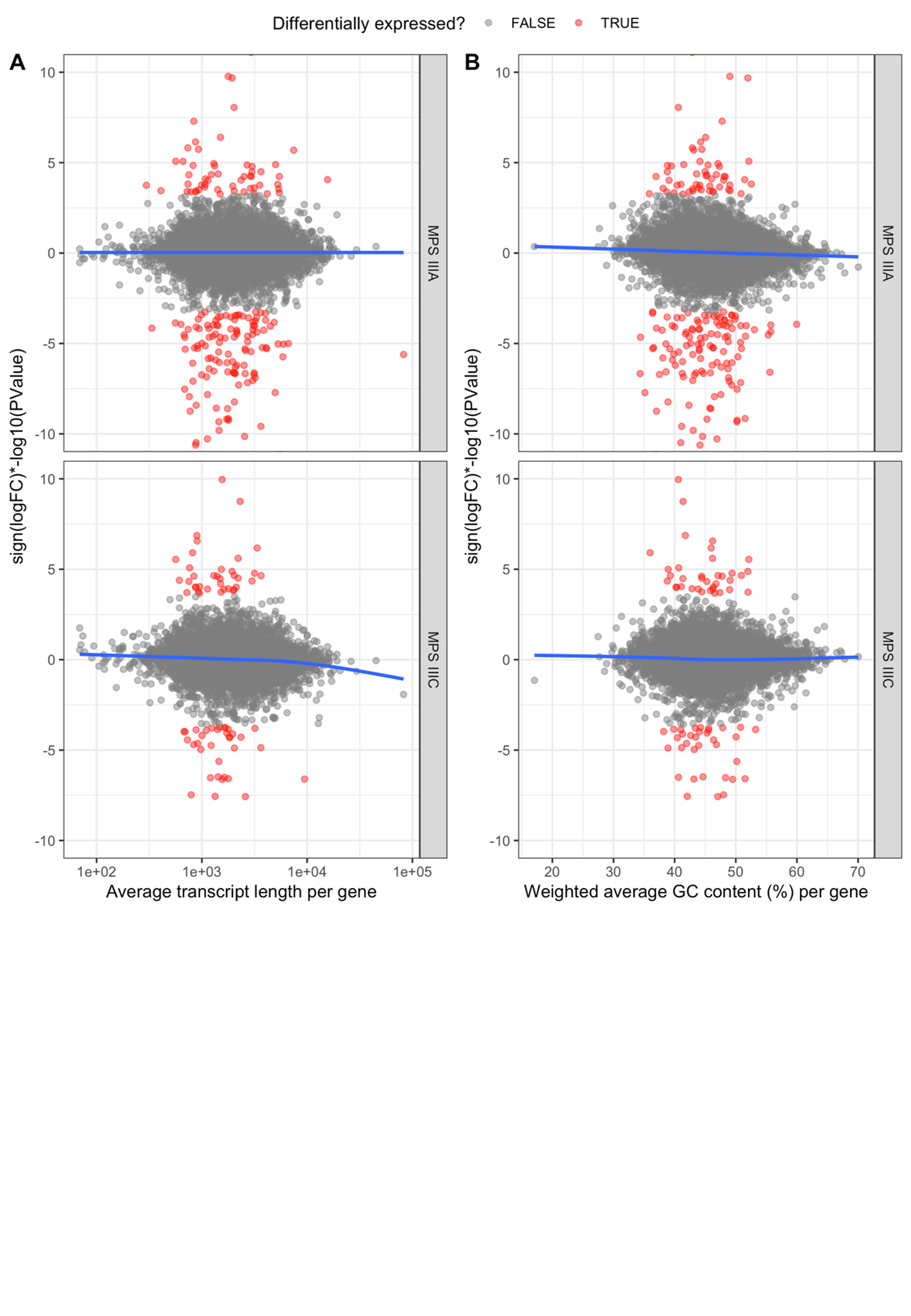

**Fig.S12:** **Observed bias for differential expression with GC content and length in the MPS IIIA and MPS IIIC cohort**. A ranking metric was calculated using the sign of logFC multiplied by − log_10_ of the p-value against **A,** the weighted (by transcript length) %GC content per gene; and **B,** the average transcript length per gene. The blue curve indicates the line of best fit from a generalised additive model. Given the gam fit is a not completely horizontal line mostly overlapping y = 0, a small bias for %GC content or length is likely present in this dataset. The ranking statistic limits were constrained to − 10 and 10 for visualisation purposes.

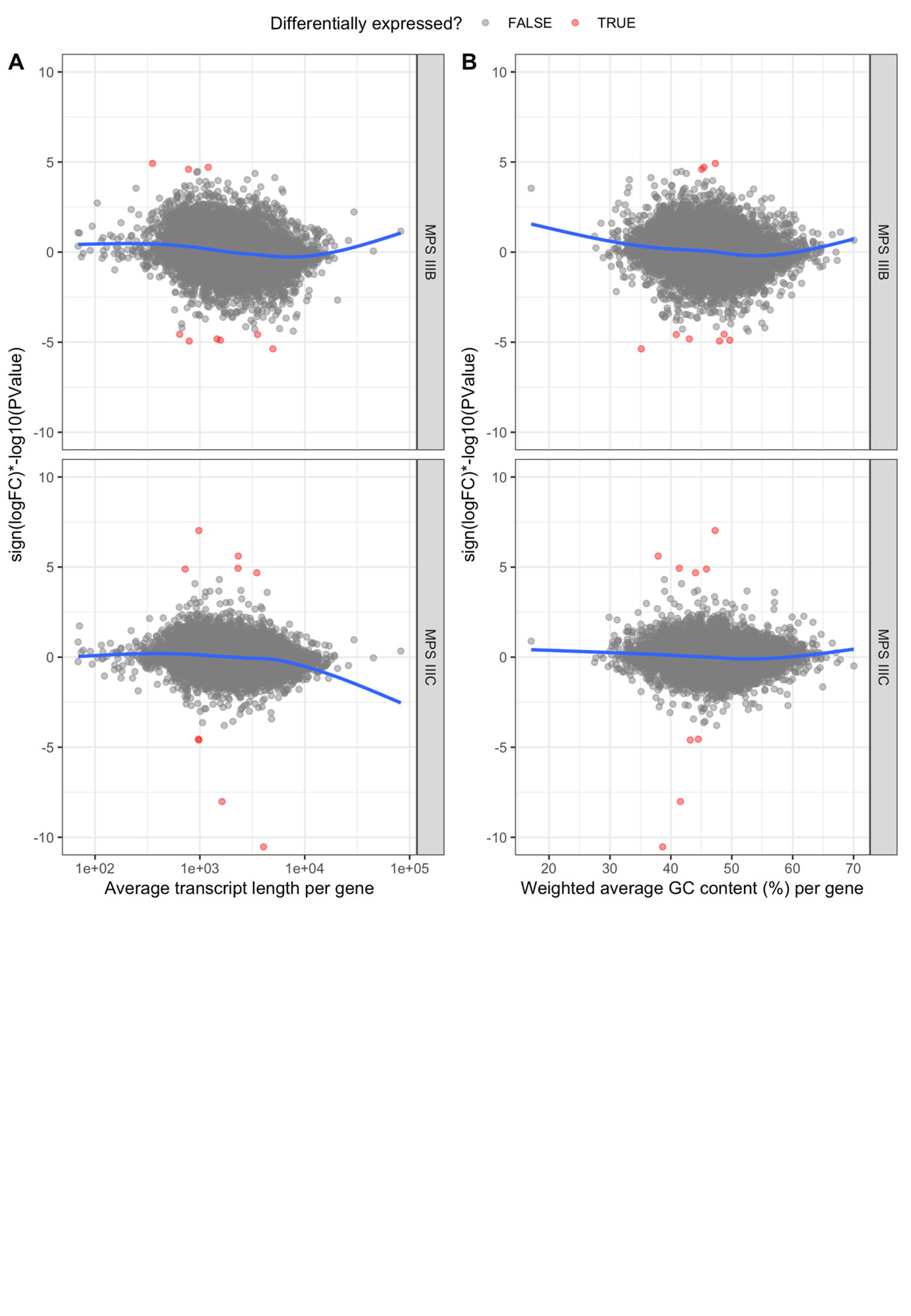

**Fig.S13:** **Observed bias for differential expression with GC content and length in the MPS IIIB and MPS IIIC cohort**. A ranking metric was calculated using the sign of logFC multiplied by − log_10_ of the p-value against **A,** the weighted (by transcript length) %GC content per gene; and **B,** the average transcript length per gene. The blue curve indicates the line of best fit from a generalised additive model. Given the gam fit is a not completely horizontal line mostly overlapping y = 0, a small bias for %GC content or length is likely present in this dataset. The ranking statistic limits were constrained to − 10 and 10 for visualisation purposes.

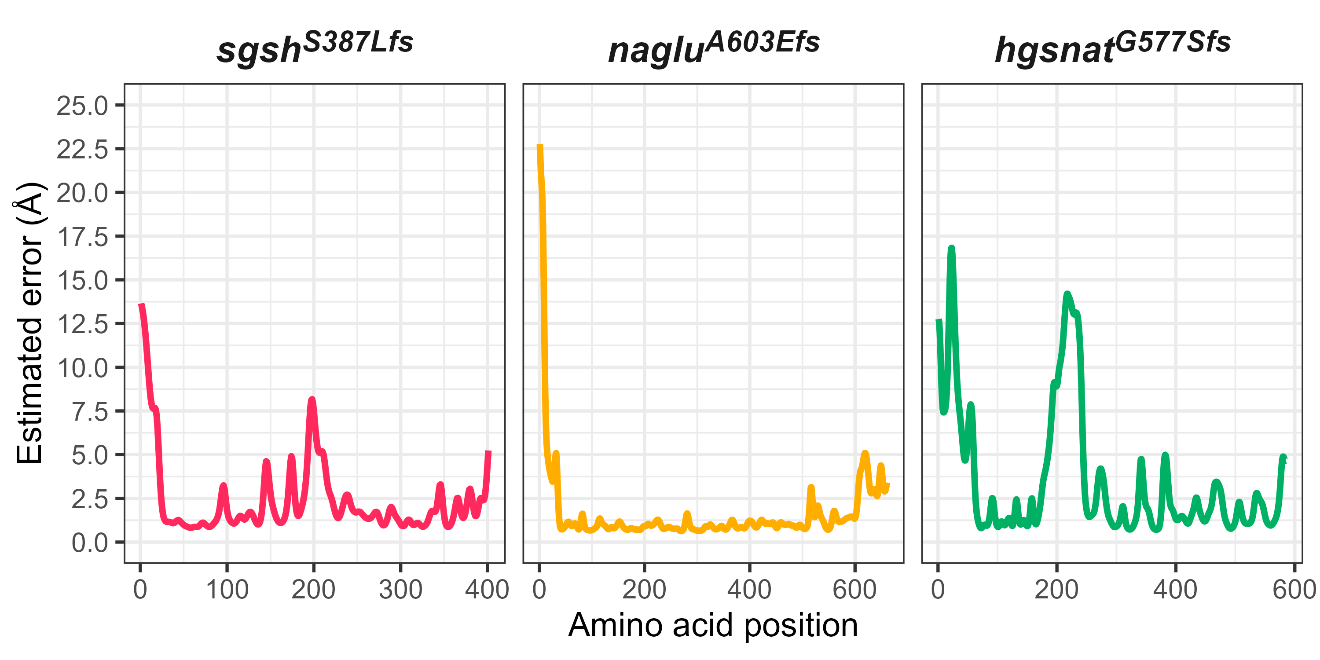

**Fig.S14: Estimated error of *sgsh^S387Lfs^* (left), *naglu^A603Efs^* (centre) and *hgsnat^G577Sfs^* (right) RoseTTAFold protein structure predictions across each amino acid residue**. Estimated error is reported in angstroms.

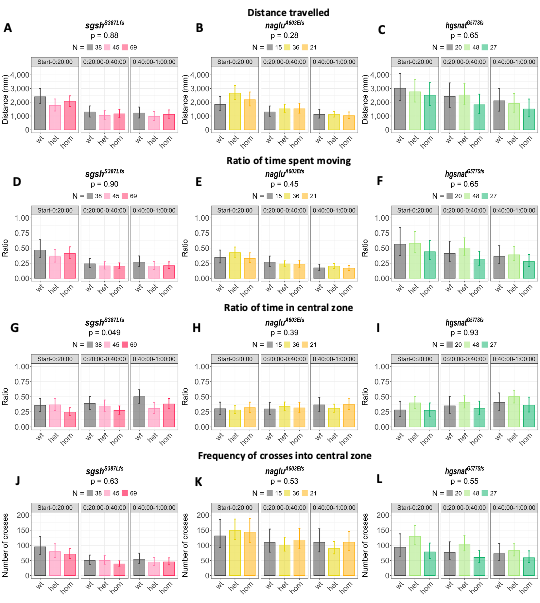

**Fig.S15: MPS III zebrafish larvae do not show behaviour phenotypes reminiscent of the human disease at 5 days post fertilisation. A-C**, Linear mixed model-predicted estimated marginal means (EMM) for total distance travelled during 1 hour (split across three 20 minute time bins) by MPS III larvae. Families consisted of larvae wild type (wt), heterozygous (het) and homozygous (hom) for each MPS III subtype indicated in the panel title. The number (N) of larvae per genotype are also indicated in the legend. **D-F**, Generalised linear mixed effect model-predicted EMM ratio of time spent moving during 1 hour by MPS III larvae, where a greater ratio represents a greater proportion of time spent moving. **G-I**, Generalised linear mixed model-predicted EMM ratio of time in the central zone, where a greater ratio represents a greater proportion of time in the centre. **J-L**, Generalised linear mixed model-predicted mean frequency of crossings into central zone. Raw data points are shown in **Fig.S16** below.

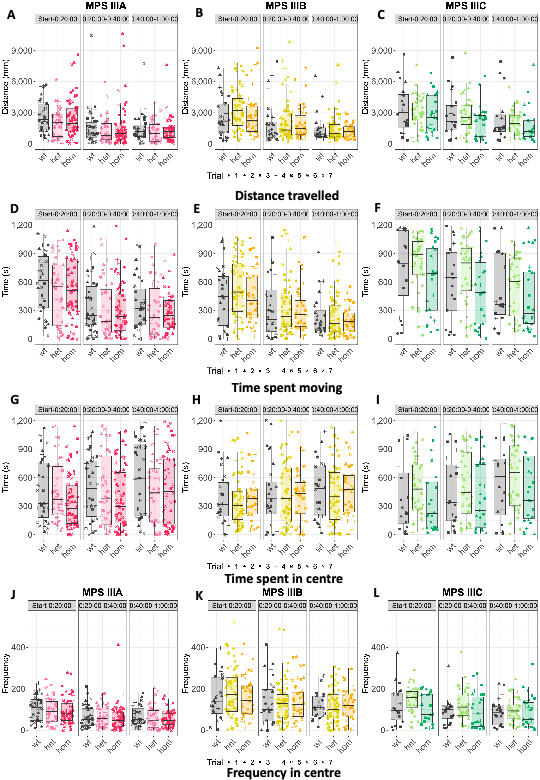

**Fig.S16: Raw data of behaviour exhibited by zebrafish larval families** at 5 days post fertilisation during a 1 hour open field test, split into 3 consecutive time bins. Families consisted of individuals wild type (wt), heterozygous (het) and homozygous (hom) for the *sgsh^S387Lfs^* (**A, D, G** and **J**)*, naglu^A603Efs^* (**B, E, H** and **K**) or *hgsnat^G577Sfs^* (**C, F, I** and **L**) mutations. Measurements of behaviour include distance travelled (**A-C**), time spent moving (**D-F**), amount of time in the central zone (**G-I**) and frequency of crossings into the central zone (**J-L**).

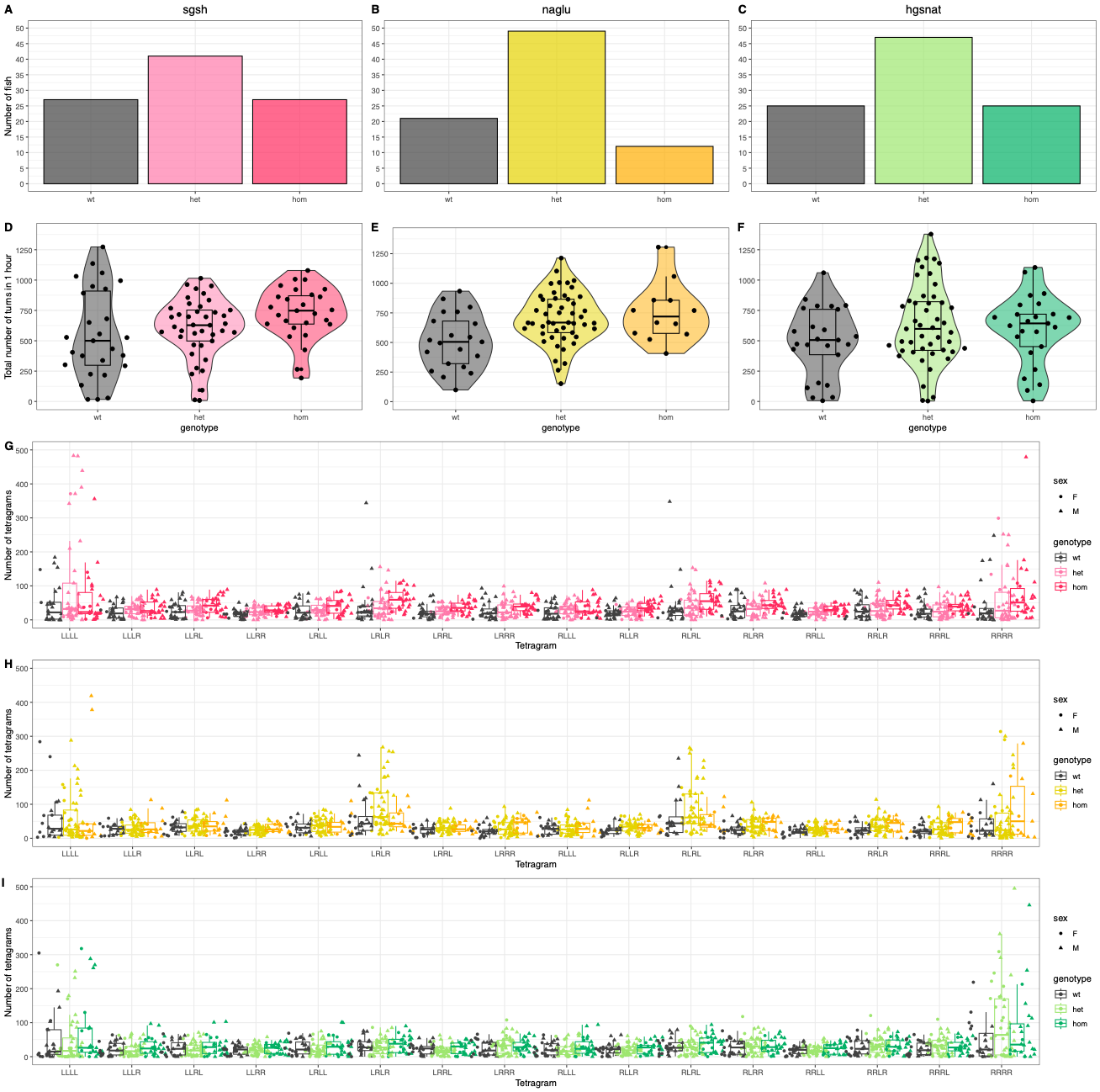

**Fig.S17: A-C,** Genotype proportions of MPS III mutant zebrafish used for behavioural testing in the Y-maze. Counts of fish wild type, heterozygous and homozygous for **A,** *sgsh^S387Lfs^* **B**, *naglu^A603Efs^* and **C**,*hgsnat^G577Sfs^* are shown. **D-F,** violin plots of total turns performed by MPS III zebrafish during 1 hour in the Y-maze. Each individual fish is represented by a raw data point, with boxplots overlaid indicating 5 number summaries. **G-I** boxplots of number of tetragrams performed per fish during 1 hour in the Y-maze. Each individual fish is represented by a raw data point, with boxplots overlaid indicating 5 number summaries.

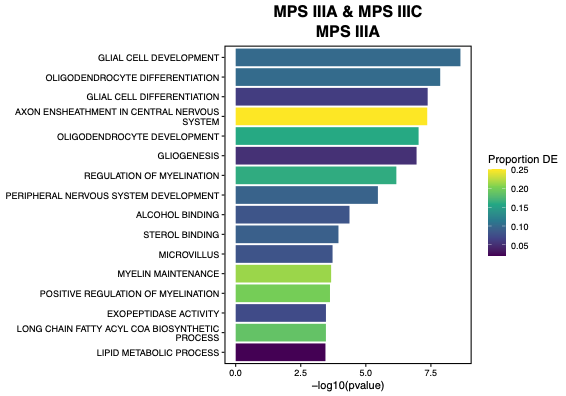

**Fig.S18: Significantly over-represented GO terms in MPS IIIA (homozygous mutant) zebrafish brain transcriptomes in the MPS IIIA and MPS IIIC cohort.** Only GO terms which reached the threshold of FDR-adjusted p-value < 0.05 are displayed.

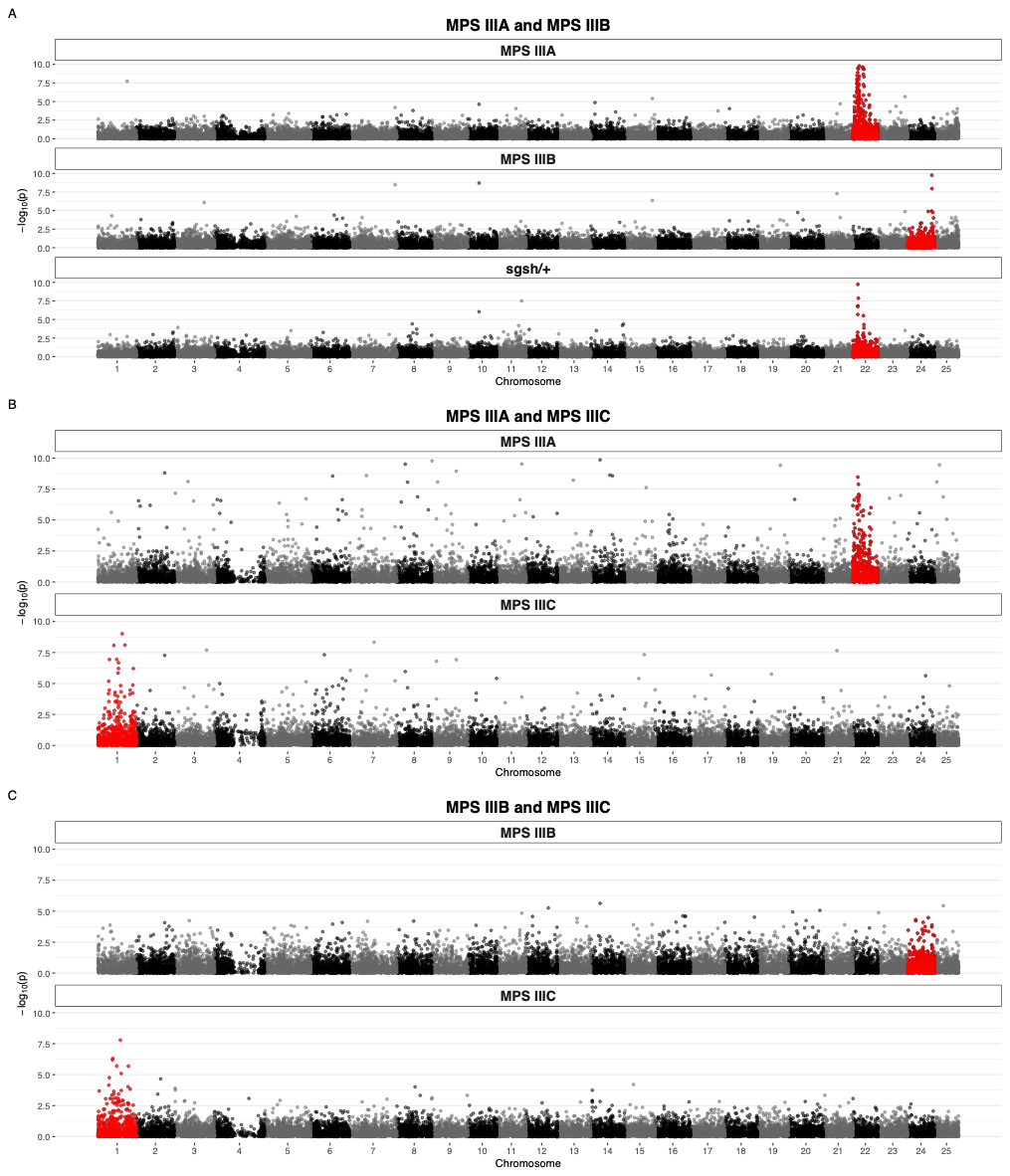

**Fig.S19:** Manhattan plots indicating p-values from the differential gene expression analyses in the **A,** MPS IIIA and MPS IIIB cohort; **B,** MPS IIIA and MPS IIIC cohort; and **C,** MPS IIIB and MPS IIIC cohort. Points appear red if they are located on the chromosome bearing the mutation.

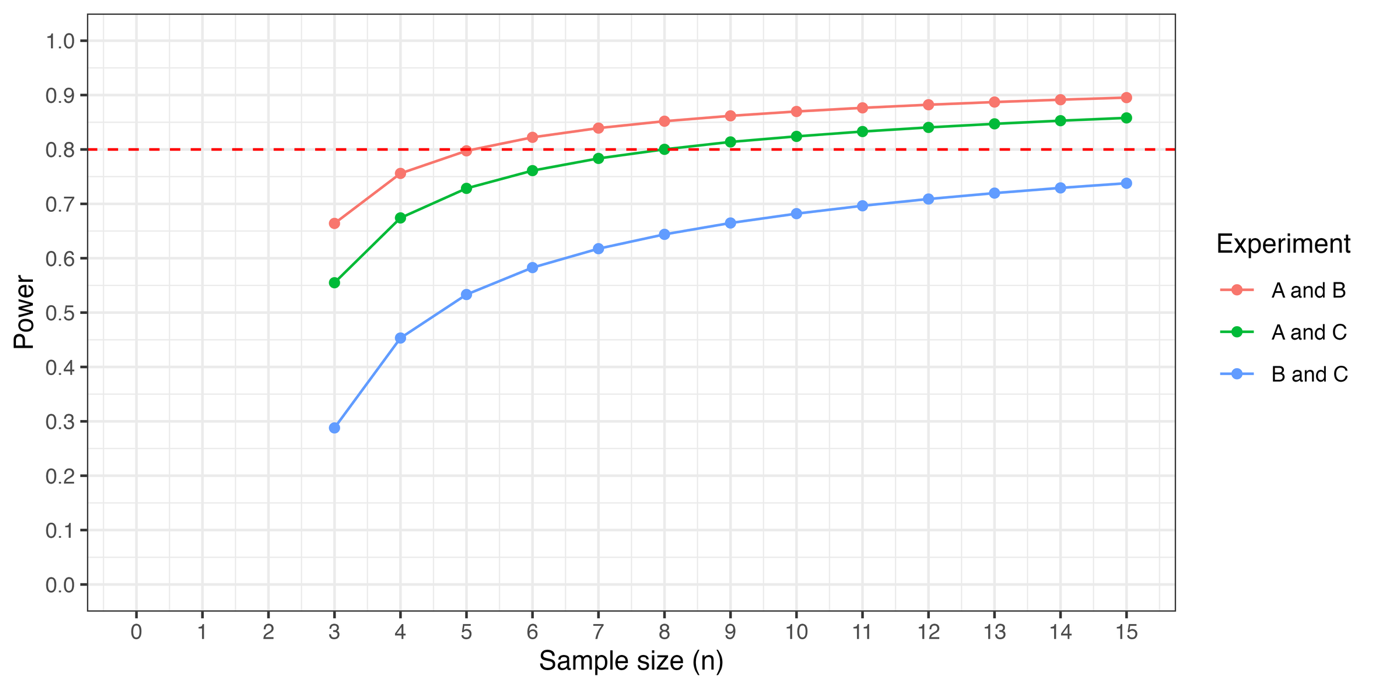

**Fig.S20:** *Post-hoc* power curves as calculated using ssizeRNA.

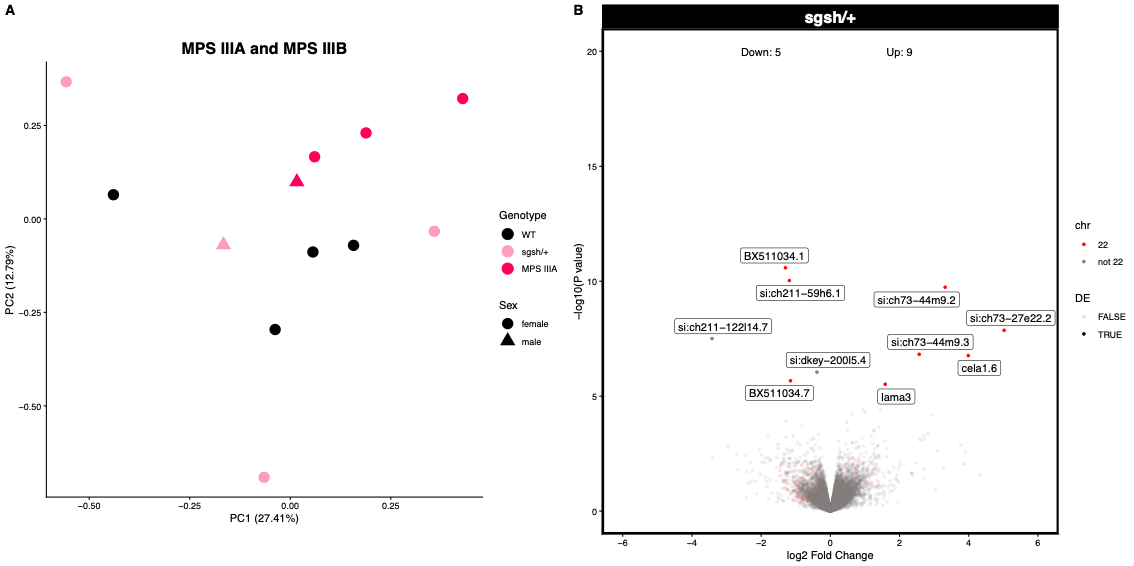

**Fig.S21: No evidence that heterozygosity for a mutation in *sgsh* affects the brain transcriptome. A,** Principal component analysis on the MPS IIIA and MPS IIIB cohort omitting the MPS IIIB brains. Each point represents a zebrafish brain transcriptome which is coloured according to its *sgsh* genotype: wild type (WT), *sgsh* heterozygous (sgsh/+) or *sgsh* homozygous (MPS IIIA). **B,** Volcano plot of differential gene expression analysis for *sgsh* heterozygous carriers. Plots are constrained between -6 and 6 on the x axis and between 0 and 20 on the y axis for visualisation purposes. Genes are indicated by points which are coloured red if they are located on chromosome 22 and labelled if the FDR-adjusted p-value was less than 0.05.
